# Ecological theory predicts ecosystem stressor interactions in freshwater ecosystems, but highlights the strengths and weaknesses of the additive null model

**DOI:** 10.1101/2020.08.10.243972

**Authors:** Benjamin J. Burgess, Drew Purves, Georgina Mace, David J. Murrell

## Abstract

Understanding and predicting how multiple co-occurring environmental stressors combine to affect biodiversity and ecosystem services is an on-going grand challenge for ecology. So far progress has been made through accumulating large numbers of smaller-scale individual studies that are then investigated by meta-analyses to look for general patterns. In particular there has been an interest in checking for so-called ecological surprises where stressors interact in a synergistic manner. Recent reviews suggest that such synergisms do not dominate, but few other generalities have emerged. This lack of general prediction and understanding may be due in part to a dearth of ecological theory that can generate clear hypotheses and predictions to tested against empirical data. Here we close this gap by analysing food web models based upon classical ecological theory and comparing their predictions to a large (546 interactions) dataset for the effects of pairs of stressors on freshwater communities, using trophic- and population-level metrics of abundance, density, and biomass as responses. We find excellent overall agreement between the stochastic version of our models and the experimental data, and both conclude additive stressor interactions are the most frequent, but that meta-analyses report antagonistic summary interaction classes. Additionally, we show that the statistical tests used to classify the interactions are very sensitive to sampling variation. It is therefore likely that current weak sampling and low sample sizes are masking many non-additive stressor interactions, which our theory predicts to dominate when sampling variation is removed. This leads us to suspect ecological surprises may be more common than currently reported. Our results highlight the value of developing theory in tandem with empirical tests, and the need to examine the robustness of statistical machinery, especially the widely-used null models, before we can draw strong conclusions about how environmental drivers combine.

## Introduction

Ecosystems are being subjected to a wide variety of external stressors (Halpern et al. 2015), acting across terrestrial, freshwater, and marine biomes (Scheffers et al. 2016). Stressors, also termed drivers, factors, or perturbations (Orr et al. 2020), are frequently anthropogenic in origin (Vörösmarty et al. 2010; Geldmann et al. 2014), but are capable of being abiotic or biotic (Przeslawski et al. 2015), and are able to act at any scale, from local to global (Ban et al. 2014; França et al. 2020). While individual stressors, (e.g. climate change, habitat alteration, or pollution), are themselves capable of inducing changes in biodiversity or ecosystems and their services (Dirzo et al. 2014; Tittensor et al. 2014; Newbold et al. 2015), ecosystems are frequently, if not predominately, acted upon by multiple stressors simultaneously (Crain et al. 2008). Despite the negative connotations surrounding the term *stressor*, stressors are capable of inducing effects that are either beneficial or detrimental to the affected ecosystem (Kroeker et al. 2017). Accordingly, one of the grand challenges facing ecologists is to be able to predict and understand how these different types of ecosystem stressors interact to affect biodiversity and ecosystem services (Hodgson & Halpern 2018); though these interactions can be challenging to predict as the observed interactions can substantially deviate from what is anticipated (Christensen et al. 2006). Ultimately, knowledge of how stressors interact is important in guiding conservation and management initiatives, and in helping to prevent remediation measures from being ineffective, or even potentially harming those systems they are intended to preserve (Brown et al. 2013; Côté et al. 2016).

Aquatic ecosystems and communities are particularly threatened by multiple stressors (Dirk et al. 2020); for instance, Halpern et al., (2008) describe how every marine area is subjected to human influence, with 41% of these areas being impacted by multiple stressors. Moreover, freshwaters represent some of the most at-risk ecosystems and are frequently exposed to a wide range of stressors (Hecky et al., 2010; Ormerod et al. 2010; Woodward et al., 2010; He et al., 2019), with freshwater biodiversity declining at rates exceeding even those of the most impacted terrestrial ecosystems (Sala et al., 2000), and potentially endangering vital ecosystem services (Malaj et al. 2014). While stressors often interact to impact freshwater ecosystems (Dirk et al. 2020), their presence in freshwater systems is not a new phenomenon, with some freshwater bodies having been subjected to stressors for several centuries (Dudgeon et al., 2006). However, the stressors that freshwater systems are currently facing has expanded, with the introduction of novel stressors, such as nanomaterials, while existing stressors are continuing to have severe impacts (Reid et al., 2019). Similarly, the cumulative impact of multiple stressors has been identified as one of the most pressing and emerging threats to freshwater biodiversity, but despite this, our current understanding of both how stressors interact, and the severity of their effects, is poor (Reid et al., 2019).

The term *ecological surprise* (*sensu* Paine et al. 1998) is often used to describe the changes in a variable that contrast those anticipated when multiple stressors interact (e.g. Christensen et al. 2006; Jackson et al. 2016). Most often, the term is applied to the interactions of stressors which interact synergistically; in other words, the observed change in a variable is greater than expected under the assumption the interaction is equal to the sum of the independent stressor effects. Accordingly, the synergistic interactions of multiple stressors are important to document, firstly due to their potential to have a dramatic effect on ecological communities, and secondly because the presence of a synergistic interaction means management strategies can potentially have a large effect by mitigating against just one of the interacting stressors (Brown et al. 2013; Côté et al. 2016; Haller-Bull & Bode 2019). Because of their potential impact there has been a great deal of effort in documenting the frequency of synergy in stressors across different ecosystems and communities (Côté et al. 2016). However, there is always a danger that an emphasis on their importance could lead to overestimating the frequency of ecological surprises within the multiple stressor literature and, as highlighted by Côté et al. (2016), the evidence that most stressors interact in a synergistic manner is far from compelling. A pertinent question which has yet to be fully answered is whether synergistic interactions, or other forms of ecological surprise, really should be expected, or whether the prevalence of these interactions are skewed in some way by reporting biases, statistical sampling, or both.

However, there is relatively little ecological theory that predicts when and how often the cumulative effects of pairs of stressors should be synergistic, or indeed any other type of interaction. This is in contrast to other ecological interactions, such as the effects of multiple predators on prey density and biomass, where a much richer body of theory that has been able to generate a number of hypotheses for testing (Sih et al. 1998; Schmitz 2007). Instead, progress on ecosystem stressor interactions has been made largely by meta-analyses across a number of experiments, realms, trophic levels, measured traits, taxonomic groups, and stressor types (e.g. Crain et al. 2008; Darling and Côté 2008; Wu et al. 2011; Przeslawski et al. 2015; Jackson et al. 2016). Within ecological research, the most popular approach is to use the additive null model where the stressor interaction is predicted to be simply the sum of their individual effects (e.g. Crain et al. 2008; Darling & Côté 2008; Strain et al. 2014; Jackson et al. 2016), though the multiplicative null model is also relatively common (e.g. Bancroft et al. 2008; Gruner et al. 2008; Harvey et al. 2013; Rosenblatt & Schmitz 2014). Predominately, these null models classify interactions as either being null (the simplest additive or multiplicative effect of interacting stressors), synergisms, or antagonisms (i.e. the effect of the interacting stressors is less than expected). While distinctions are increasingly being made for various forms of antagonistic interactions (e.g. Jackson et al. 2016), there exists a range of other classification schemes (Orr et al. 2020), implemented across a number of studies (e.g. Travers-Trolet et al. 2014; Piggott et al. 2015a). This can make it difficult to generalise results across different studies, because a ‘synergistic’ or ‘antagonistic’ interaction may have contrasting definitions depending on the scheme being used. Despite meta-analyses being a powerful tool for investigating multiple stressors, they have to date highlighted no general covariates capable of explaining of the broad patterns of multiple stressor interactions, which in turn lead to more general predictions of the consequences of multiple stressors (Côté et al. 2016).

Given the lack of consistent generalities from empirical studies, there have been calls for the development of theory within multiple stressor research. Of primary interest is the generation of theory which can provide a mechanistic underpinning to the field, and hopefully allow for better prediction and an increased understanding of multiple stressor interactions, compared to that which is provided solely by a null model approach (De Laender 2018). For example, using only statistical null models it is hard to predict, and therefore understand, how an interaction between stressors will change as one or more stressors changes in intensity. Some theory has been developed for particular case studies (e.g. Brown et al. 2013; Galic et al. 2018), but only a few studies have so far looked for more general insights. For example, Haller-Bull and Bode (2019) used three population dynamic models to investigate how stressors reducing population growth or supressing carry capacity combine to affect equilibrium population biomass under harvesting. Across all models they found synergy only occurs if there are several impacts on growth rate, and more generally the interaction behaviour can be predicted by the relationship between the impacted parameter and the equilibrium population size; a convex relationship implies antagonism, and a concave relationship implying synergy.

Although population models are easier to analyse, incorporating trophic interactions would seem a necessary feature for a general dynamical theory for multiple ecosystem stressors since they may act either directly (e.g. on mortality rate of a given species) or indirectly (e.g. on mortality rate of the prey of a given species). Indeed, De Laender (2018) has recently argued for the use of resource uptake theory to make predictions about stressor interactions, and, as an example, showed that in a two-species community, the manner in which stressors interact is dependent on the details of which species (one, or both) are being directly affected by the stressors. Extending to more diverse ecological communities, Thompson et al. (2018a) used modified (log-linear) Lotka-Volterra models to investigate how the effect of multiple stressors on species richness changes with the type of biological interaction that dominates a community. They found negative biological interactions, (predation, competition), were more likely to lead to synergistic changes in species richness, whilst stressor interactions were predominantly additive or slightly antagonistic when biological interactions were positive. These models all show much promise for theory to generate predictions, but as yet none have been tested against data. To compare to data, models need to incorporate stochasticity to mirror the sampling variation found in the real world. In natural experiments sampling variation occurs in the estimation of the state variables of interest such population density or biomass (e.g. Graham & Vinebrooke 2009; Piggott et al. 2015b), or individual growth rates (e.g. Reisinger & Lodge 2016), and this error enters the estimation of the interaction of the co-occurring stressors with the inevitable result that some interactions are misclassified due to sampling variation. The simplest way to incorporate sampling variation in models is via some form of observation error, but De Laender (2018), Haller-Bull and Bode (2019), and Thompson et al. (2018a) all base their predictions on deterministic models, with stochasticity only entering the latter in the form of parameter combinations.

Here we build on this theory by developing classical community ecology models based upon Lotka-Volterra consumer-resource dynamics, but including observation error, to generate predictions from biologically simple food webs. These predictions are tested against an extensive dataset for the effects of co-occurring stressor interactions on the biomasses and densities of freshwater organisms, taken from a review of the experimental literature. Using this twin approach, we answer the following questions: Can dynamical food web theory predict (1) the frequencies of stressor interaction types across the individual experimental studies, and/or (2) the expected summary effect sizes and summary interaction type in a meta-analytical framework? In particular we ask if the apparent absence of ecological surprises in the empirical literature is expected from ecological theory, and in so doing we also test the robustness of the currently popular additive null model for classifying stressor interactions to sampling variation. As will be shown below, our results uncover a high level of agreement between theory and data but also highlight some of the strengths and weaknesses of the additive null model.

## Materials and Methods

### Theoretical Models

In order to provide a theoretical underpinning for the empirical results, we build food chain models using the classical Lotka-Volterra consumer resource equations. To increase the robustness of our conclusions we consider two forms of model; one where (within trophic level) density dependence affects the death rates of each trophic level, and a second where consumer uptake is density regulated (Table 1). Both these scenarios were analysed by Heath et al., (2014) to investigate the roles of different types of density dependence on trophic cascades, and more detail can be found there. In both models the basal level of the chain describes dynamics of a key nutrient that limits the productivity of the food chain, and we assume nutrients are added at a constant rate, ω. Each subsequent equation then describes a different type of consumer. The first level is wholly dependent on the nutrients and could represent a primary producer such as an algal species that requires a key mineral such as silica. The second level consumes the first trophic level and is in turn consumed by a third trophic level, and so on until the apex consumer is reached. In the density dependence model (Equation 1, Table 1), the consumer *i* exploits the resource (trophic level *i* – 1) with a constant consumption/attack rate, α_*i*_, and the conversion efficiency parameter, *ε*_*i*_, determines the proportion of the resource consumed that is converted into new consumers. Under density dependence, the density of the consumer is self-regulated by the intraspecific density dependence parameter *λ*_*i*_, which leads to an increase in death rate as the consumer density increases. In contrast, the consumer uptake regulation model (Equation 2, Table 1), assumes the effect of increasing consumers is to slow down the consumption of the resource, perhaps due to increased inference. In this case, the parameter *v*_*i*_, determines the consumer density at which the maximum per capita uptake rate is halved, defined as the density *x*_*i*_ =1/*v*_*i*_.

**Table 1:**
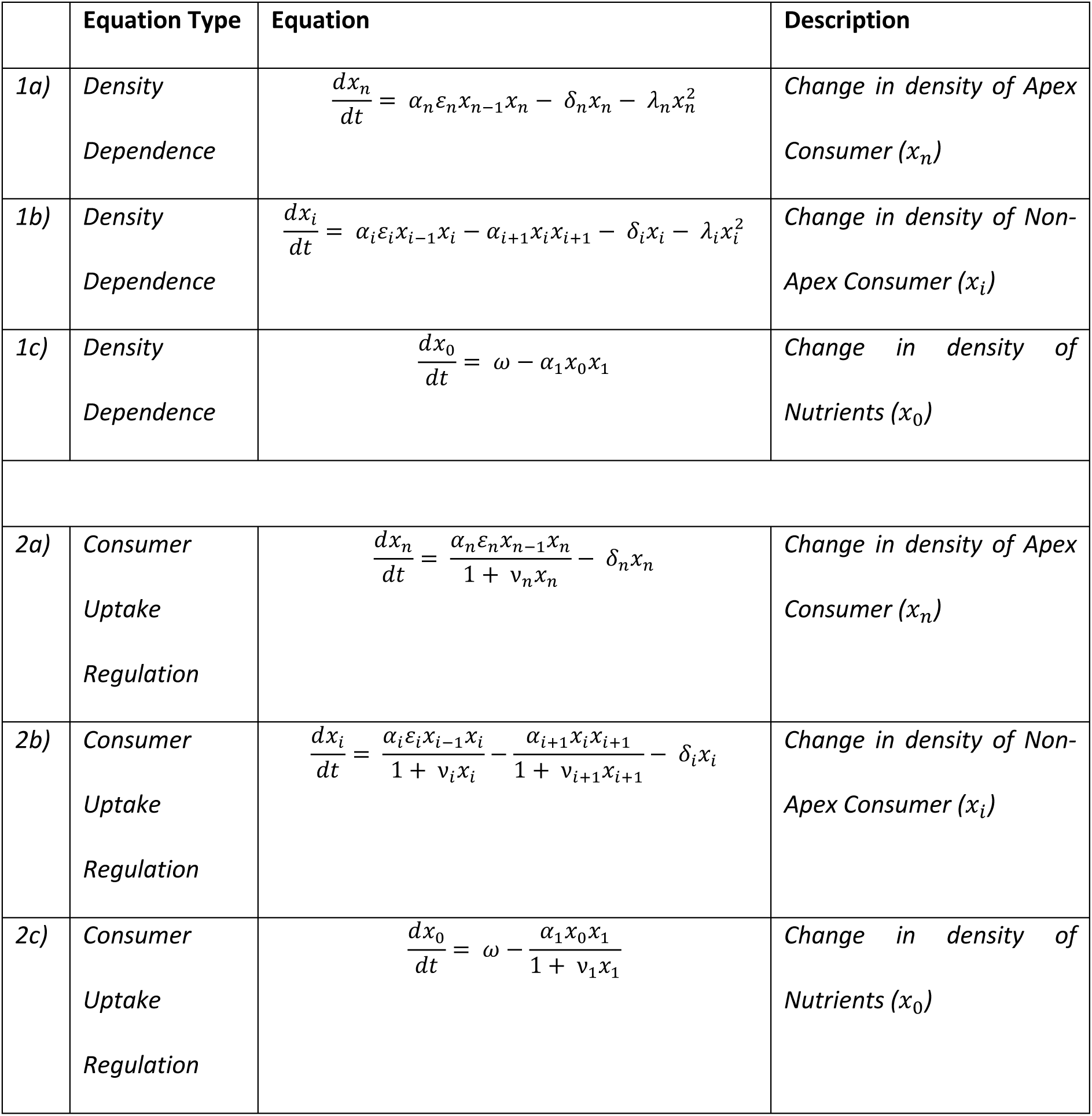
Equations used to establish theoretical food-chains. The equations, sets, and a brief description of the equivalent ecological trophic are shown.

|  | Equation Type | Equation | Description |
| --- | --- | --- | --- |
| 1a) | Density<br>Dependence | $\frac{dx_n}{dt} = \alpha_n \varepsilon_n x_{n-1} x_n - \delta_n x_n - \lambda_n x_n^2$ | Change in density of Apex<br>Consumer ($x_n$) |
| 1b) | Density<br>Dependence | $\frac{dx_i}{dt} = \alpha_i \varepsilon_i x_{i-1} x_i - \alpha_{i+1} x_i x_{i+1} - \delta_i x_i - \lambda_i x_i^2$ | Change in density of Non-Apex Consumer ($x_i$) |
| 1c) | Density<br>Dependence | $\frac{dx_0}{dt} = \omega - \alpha_1 x_0 x_1$ | Change in density of<br>Nutrients ($x_0$) |
| 2a) | Consumer<br>Uptake<br>Regulation | $\frac{dx_n}{dt} = \frac{\alpha_n \varepsilon_n x_{n-1} x_n}{1 + v_n x_n} - \delta_n x_n$ | Change in density of Apex<br>Consumer ($x_n$) |
| 2b) | Consumer<br>Uptake<br>Regulation | $\frac{dx_i}{dt} = \frac{\alpha_i \varepsilon_i x_{i-1} x_i}{1 + v_i x_i} - \frac{\alpha_{i+1} x_i x_{i+1}}{1 + v_{i+1} x_{i+1}} - \delta_i x_i$ | Change in density of Non-Apex Consumer ($x_i$) |
| 2c) | Consumer<br>Uptake<br>Regulation | $\frac{dx_0}{dt} = \omega - \frac{\alpha_1 x_0 x_1}{1 + v_1 x_1}$ | Change in density of<br>Nutrients ($x_0$) |

Using these equations, we establish food-chains comprising either three, four, or five trophic levels, and the equation for each trophic level models how the biomass or density changes over time. For simplicity we assume all key parameters (nutrient input ω; consumption rates α_*i*_; conversion efficiencies *ε*_*i*_; uptake regulators *v*_*i*_; density independent δ_*i*_, and dependent death rates *λ*_*i*_, for trophic level *i*) do not vary over time, and we investigate the effect of stressors on equilibrium biomasses/densities. The models do not consider any spatial structure in the community which also remains closed to immigration from outside apart from the constant input of the nutrient. Hence these models represent the simplest form of community dynamics that could be used to investigate the effects of multiple stressors and how they interact.

Stressors to the food chains are modelled by changing the values for parameters and comparing the resultant equilibrium densities/biomasses across all trophic levels to the equilibria for a set of baseline parameter values. Equations 1 and 2 are not mechanistic models for specific stressors, (e.g. pollution, temperature), but instead capture the net effect of stressors on the vital rates of the food web species. For simplicity, we assume each stressor has either a positive or negative effect on one vital rate, (i.e. model parameter), and we investigate how pairs of stressors interact to affect community densities. Both the baseline parameters and the parameters after perturbation are drawn from uniform distributions with ranges given in Table 2. So, for a given food chain the baseline parameters for all trophic levels are independently sampled from the distribution of values given in Table 2. The vital rate affected by each stressor is randomly selected from the possible candidates, and the intensity of its effect on the baseline rate is drawn from a uniform distribution with ranges given in Table 2. The baseline parameter set therefore represents the control community, and as in experimental studies (e.g. Matthaei et al., 2010; Davis et al., 2018) we manipulate our model communities by investigating the effect of each stressor acting alone, as well as the stressors acting in combination. From these cases we then compute the type of stressor interaction and how they combine to the alter the community biomasses (see below for definitions of how stressor interactions are computed). To do this we choose one trophic level at random from the entire food chain but excluding the nutrient level. We focus on this population/trophic level and mirror it in our selection of empirical data (see below). This also means the species or trophic levels under scrutiny are not always directly affected by the stressor but could be affected solely due to a trophic cascade effect. It is also important to note that a stressor could lead to either an increase or a decrease in parameter value relative to the baseline; and that multiple stressors could act on the same, or different trophic level, but that each stressor affects only one parameter (and therefore biological process).

**Table 2:**
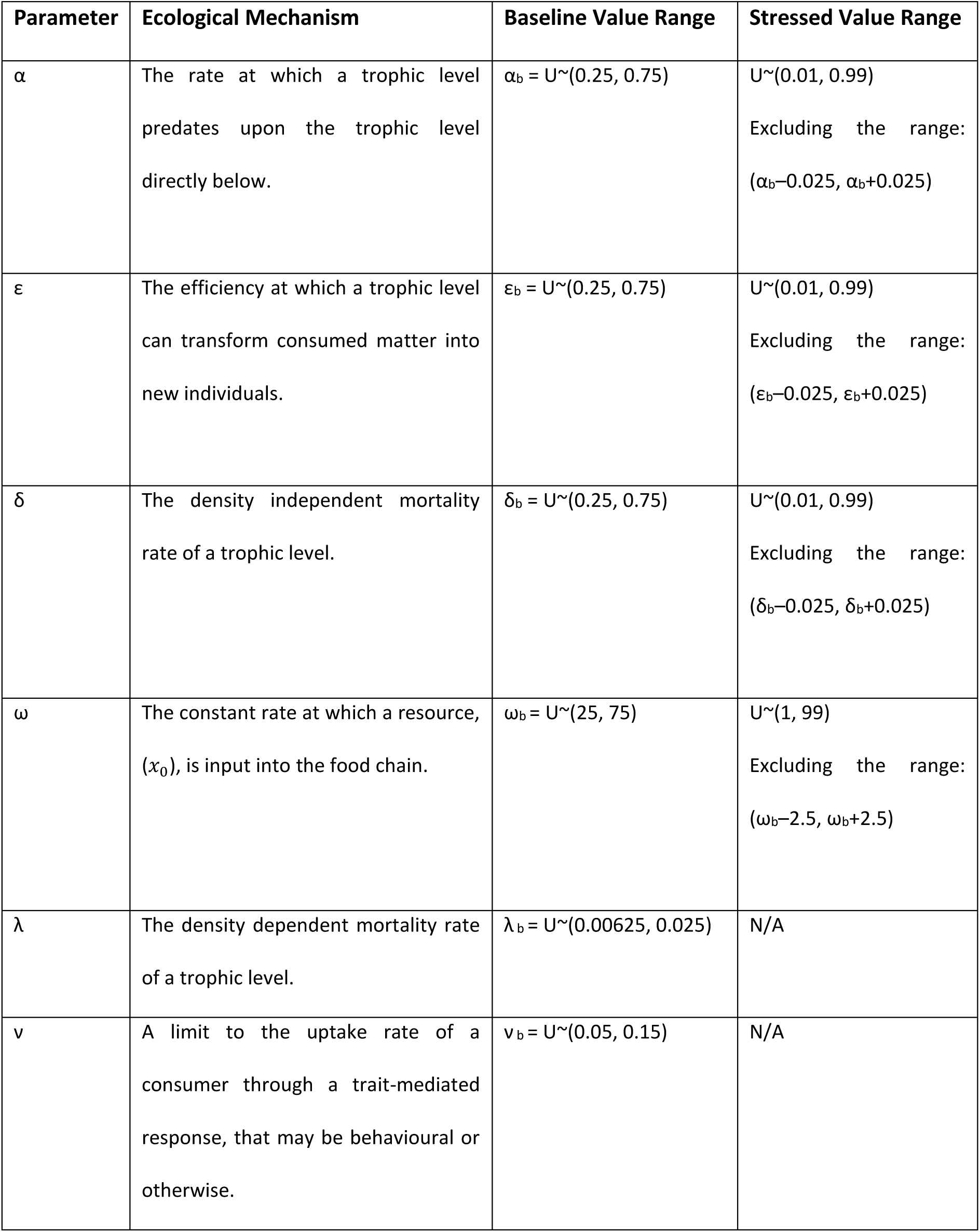
Explanation of the different parameters within Equations 1 and 2, with the mechanism they reflect, alongside the minimum and maximum values for the ranges of baseline and stressed parameter values. Each parameter is drawn from a uniform distribution U∼(a, b) with lower limit, a, and upper limit, b.

Overall, 1,320,000 different combinations, of equations, food-chain lengths, stressors pairs, and randomly selected baseline values were generated. Equilibrium densities, for each of these combinations, were calculated using Mathematica 10.4, (Wolfram Research, Inc., 2016), with equilibria and stability analyses as given in Heath et al., (2014); (for more details see Supplementary Material 1). We only consider cases where the equilibria are all stable, and feasible (i.e. all densities were positive), and only equilibrium densities for trophic levels *x*_*1*_ and above are included in the stressor interaction results i.e. we exclude the nutrient level from our stressor interaction analyses. Across all 1,320,000 combinations, 79.9% of the parameter sets result in the determination of equilibrium densities that are both stable and feasible, with the discarded 20.1% parameter sets resulting in at least one biologically unfeasible density/biomass. From the full set of stable and feasible communities we select at random 360,000, and for each one randomly select a single trophic level for the focus of our estimation of the stressor interaction. All subsequent analyses of the theoretical data are performed on this group of 360,000 theoretical interactions. This subsetting is required as there is a negative relationship between number of trophic levels and likelihood of the community being both stable and feasible, which biases the full dataset towards communities with only three trophic levels. The final 360,000 stressor interactions are selected with weighted probabilities to ensure approximately one third (i.e. ∼120,000) are from each of the three food chain lengths, and that each model (Table 1) is also approximately equally represented.

Unlike the empirical studies used in the meta-analyses below, the food chain models are purely deterministic, meaning there were no random fluctuations around the equilibrium densities. In effect, for any given pair of stressors, there is no sampling error in the theoretical data. Clearly, this differs from the empirical data where sampling error leads to an estimate of the densities/biomasses under investigation in the control and treatment replicate communities, and this sampling variation may lead to some stressor interactions being misclassified. For a better comparison to the empirical data, and to test the robustness of the additive null model to sampling variation, we modelled observation (or measurement) error by taking the 360,000 theoretical interactions at equilibrium from our original analyses and then multiplying the biomass of each trophic level by a random number drawn from a Gaussian distribution with mean 1 and standard deviation *σ*. This process was repeated between three and six times for each treatment, analogous to the number of replicates per treatment found in our empirical data (see below). Thus, larger values for *σ* lead to larger deviations around the equilibrium biomasses, and therefore a larger observation error, with an increased likelihood that the stressor interaction is misclassified. Standard deviations, *σ*, are from one of 86 different levels, ranging from 1×10^−10^ to 0.5, in consistent logarithmic increments, (e.g. 8×10^−10^, 9×10^−10^, 1×10^−9^, 2×10^−9^, etc.). Supplementary Material 1 details a complete overview of how observation error was incorporated into the theoretical data.

### Collation of Empirical Data

Through use of Web of Science we searched the primary scientific literature, for papers published before 1^st^ January 2019, which investigated the impacts of multiple stressors upon freshwater communities. In order to be incorporated, papers needed to report results where there was a factorial design, namely; (i) a control (without stressors), (ii) each stressor acting individually, (iii) the stressors acting simultaneously. Papers needed to report the mean value of the response, number of replicates, and standard deviation or standard error for each treatment in the factorial design; failure to report any of this information led to the study being excluded from our analysis. Additionally, papers were required to report at least one of the following untransformed metrics: biomass, abundance, density, or chlorophyll-a of one or more groups of organisms within the stressed community. Hence, and in line with our trophic models, the focus of our effort is directed towards studies that report the effects of stressors acting at the population and community levels. Papers often report the impacts of stressors upon multiple different groups of organisms within a community; and for these the responses of all different groups of organisms were included within the overall dataset. The different groups of organisms could comprise: populations of a single species, (e.g. *Daphnia pulex*); a group of organisms within the same feeding guild, (e.g. detritivores); a group of taxonomically similar organisms, (e.g. *Ephemeroptera, Plecoptera, and Trichoptera* taxa); or a group of similar organisms, (e.g. macroinvertebrates or algae).

To be collated within our dataset, papers had to investigate communities comprising a minimum of two different groups of organisms. Studies investigated a wide range of different stressors, though these were subsequently grouped into broader categories of stressor, such as Temperature, Contamination, and Habitat Alteration.

Previous analyses have frequently focussed upon collating data for only the greatest single intensity of a stressor (e.g. Jackson et al., 2016). In contrast, where studies reported the responses of communities to multiple intensities of different stressors, data for all of the different intensities was collated. All interactions considering the different intensities of stressors were included in the overall dataset, although covariation in data due to repeated experiments across different stressor intensities were accounted for in the final meta-analyses (see section *Meta-Analytical Models*).

Some studies report multiple different response metrics for the same group of organisms, include the same species within multiple different groups, or report data for the same experiment over multiple different time points. Accordingly, in order to reduce correlation/covariance within the overall dataset, these interactions are removed from our analyses. For instance, interactions measuring density are prioritised over abundances, which are in turn prioritised over biomasses, or measurements of chlorophyll-a respectively. Similarly, where papers reported data for interactions over multiple different time points, only the final time point is used as this best matches our equilibrium assumption for the theoretical models.

Supplementary Material 2 gives a complete overview of the different search terms used to find studies, the methodology used to determine whether the data for a study could be collated, the processes for extracting and collating the data, and the process for removing interactions to prevent covariance.

### Determining of Effect Sizes and Classification of Interactions

Across both the theoretical and empirical datasets, we use the same methodology for determining the classification of an interaction, with this being implemented through use of the effect size metric, Hedges’ *d*, (Gurevitch et al., 2000). Hedge’s *d* is frequently implemented in research investigating the impacts of multiple stressors due to its ability to estimate the standardised mean difference between the means of stressed and control samples; whilst also being unbiased by small sample sizes (Hedges & Olkin, 1985). Hedge’s *d* is calculated through a comparison of the effect of the interaction to the sum of the effects of the stressors acting individually; namely, an additive null model. In line with current methodologies, we invert the sign of the interactions when the expected effect of the additive null model is negative (Piggott et al., 2015a). Following this methodology allows for interaction effect sizes to be compared regardless of their directionality. As such, we focus on the classification of the interaction as opposed to the absolute magnitude/polarity of the effects. Supplementary Material 3 gives a complete breakdown of the equations used for calculating Hedge’s *d*.

Once Hedge’s *d* for a given interaction of stressors was calculated, we then classify the interaction into one of four types as illustrated by Figure 1 and following the convention of Jackson et al., (2016). In brief, the four interaction classifications are: (i) *Additive*, where the effect of the additive null model is statistically indistinguishable from the effect of observed interaction; (ii) *Synergistic*, where the observed interaction effect is greater than the effect of the additive null model; (iii) *Antagonistic*, where the observed interaction effect is less than the effect of the additive null model, but both effects have the same polarity; (iv) *Reversal*, where the observed interaction effect is negative but the effect of the additive null model is positive. The distinction between antagonistic and reversal interactions is relatively recent (e.g. Travers-Trolet et al., 2014; Jackson et al., 2016), with most research still using the appellation of antagonistic to refer to both antagonistic and reversal interactions (e.g. Velasco et al., 2019; Gomez Isaza et al., 2020). If Hedge’s *d* is positive the interaction is classed as synergistic. If Hedge’s *d* is negative, the interaction is classed as either an antagonistic or reversal interaction, though this can only be determined by comparing the effect of the additive null model to the observed effect (as outlined above). Each value of Hedge’s *d* has corresponding 95% confidence intervals; if these confidence intervals incorporate 0 then an interaction is deemed to be additive. The classification scheme outlined above is one of a number of possible choices (e.g. Crain et al., 2008; Jackson et al., 2016), and Supplementary Material 4 details a comparison of how these different schemes to one another.

**Figure 1:**
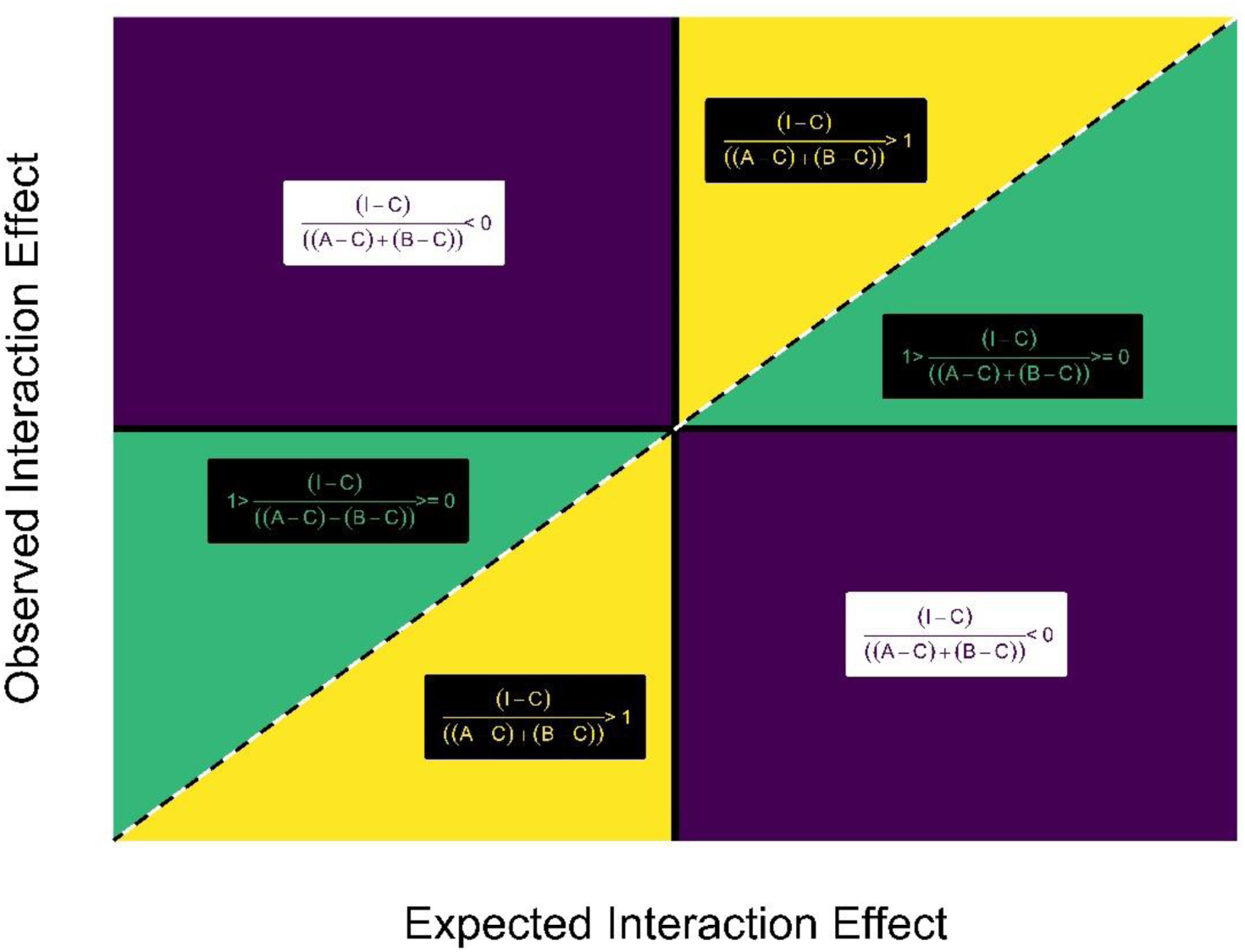
Pictorial depiction of interaction types. Additive interactions are shown by the diagonal black and white dashed line. Yellow denotes the areas occupied by synergistic interactions. Purple denotes the areas occupied by reversal interactions. Green denotes the area occupied by antagonistic interactions. Equations for the general classifications are shown for antagonistic, reversal, and synergistic interactions. C – control, A – Only Stressor A present, B – Only Stressor B present, I – Both Stressors A and B present. In order for an interaction to be classed as additive, the effect of the interaction would be equal to the sum of the effects of the individual stressors, ((I-C) = (A-C)+(B-C)).

### Vote-Counting

Following the classification of all interactions, we implement a vote-counting methodology to determine the relative proportions of the interaction classes across both the theoretical and empirical datasets. To consider the effect of different strengths of sampling variation on the ability to detect the ‘true’ stressor interaction in the modelled data, we compute the frequency of interaction types for both the case with no observation error, and for the full range of observation error levels investigated.

### Meta-Analytical Models

Alongside the vote-counting methodology, we determine the summary interactions class using a meta-analytical approach to both the theoretical and empirical datasets. The meta-analytical models are Weighted Multi-Level/Multi-Variate Random-Effect Models, and implemented in the *metafor* package (Viechtbauer, 2010) in R. For the empirical dataset random effects are specified as being the ID of the study group of organisms nested within the ID for study. The random effects are specified in order to account for both between- and within-study variation. Additionally, some studies consider multiple intensities of one or more stressors, and as such, calculations of the interaction class for each intensity of stressor use the same control. To account for any covariance between the different intensities of a single stressor, we incorporate covariance-variance matrices within the meta-analytical models. For the empirical dataset, mixed effect models are also conducted with the fixed effects of stressor pair or organism group (see Supplementary Material 5). The summary effect size for the theoretical dataset is also determined using a similar process. However, due to computational limitations caused by the number of interactions under analysis (360,000 interactions at each level of observation error), meta-analytical models for the theoretical data are fitted using the *lm* function. The models applied to both the theoretical and empirical datasets are explained in further detail within Supplementary Material 5.

The overall effect from a meta-analysis needs to be checked for consistency among effect sizes, termed as heterogeneity (Nakagawa et al., 2017). We use the *I*^2^ statistic, which is bounded between 0% and 100%, with 25%, 50%, and 75% being suggested as levels for respectively, low, medium, and high heterogeneity (Higgins et al. 2003). Ecological meta-analyses often report high levels of heterogeneity (Senior et al., 2016), perhaps due to the variation in study organisms common to the questions being asked, and we might expect a high value here due to both range of study organism and range of stressor type. To explore the potential causes of heterogeneity within the empirical meta-analysis, we conduct separate meta-analyses upon two sub-groups of the dataset, a similar process to running a meta-regression (Nakagawa et al., 2017), using organism group (i.e. producer or consumer) as the categorical moderators to explore heterogeneity (see Supplementary Material 6). We also consider publication bias (see Supplementary Material 6); though it should be noted that common tests for publication bias within meta-analyses can be limited by high heterogeneity (Nakagawa et al., 2017).

### Comparison of Theoretical and Empirical Data

Using the methods outlined above we ask whether the theoretical models are good predictors for (1) the respective frequencies of the different interaction types; and (2) the summary interaction class returned from the meta-analyses of the freshwater experimental literature on the effects of co-occurring stressors. Under the assumption that all empirical studies involve some observation (measurement) error we compare the empirical data to the model generated interactions that include observation error levels between 1×10^−2^ and 0.5 (a total of 5,040,000 modelled interactions).

## Results

### Stressor Interactions within Theoretical Data

We find no strong difference between classification of stressor interactions from either form of food chain model (Table 1), nor between the different length food chains (see Supplementary Material 1), showing the frequencies of interaction are robust to these details of the models. For the entire theoretical dataset of 360,000 interactions, (comprising both Consumer Uptake Regulation and Density Dependence Equations, and across food chains of three, four and five levels), without observation error, antagonistic and synergistic interactions are the most frequently assigned (0.483 and 0.480 respectively), followed by reversal (0.0288), and finally additive interactions (0.00856). However, these interaction frequencies are very sensitive to observation error. Increasing observation error leads to more interactions being classified as additive, (the null model), and at likely realistic levels, additive interactions are clearly dominant (Figure 2a).

**Figure 2:**
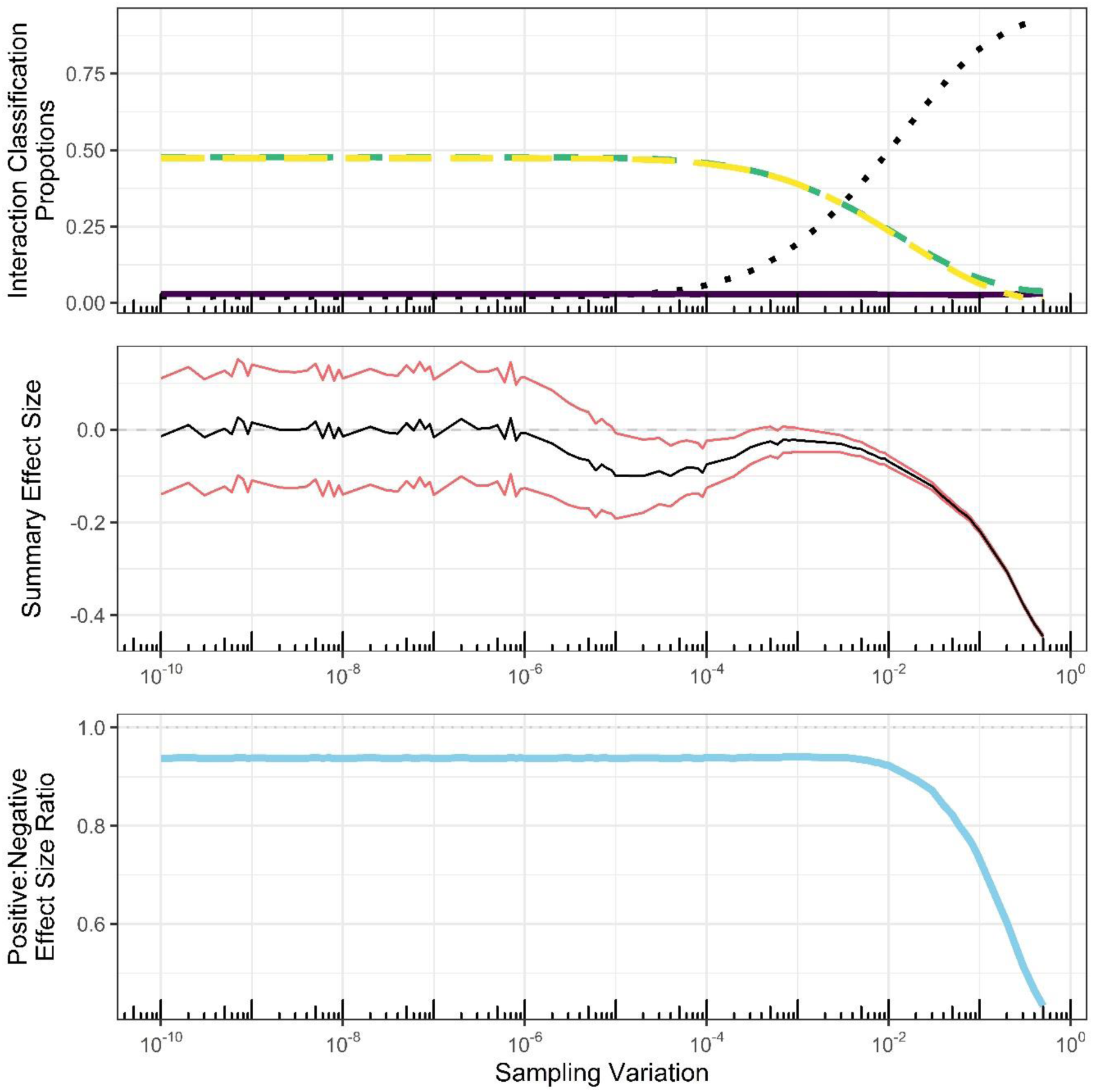
The effect of sampling variation on the stressor interaction categorisation, and summary meta-analytic effect sizes in the theoretical data. (Top Panel) Proportions of the different interaction classes for the 360,000 theoretical interactions at each level of sampling variation. Dotted black line denotes additive interactions. Green short-dashed line indicates antagonistic interactions. Yellow long-dashed line denotes synergistic interactions. Purple line indicates reversal interactions. (Middle Panel) Summary effect sizes for the 360,000 theoretical interactions, at each level of sampling variation. Black lines denote summary effect sizes, and red lines denote 95% confidence intervals. (Bottom Panel) Ratio of positive to negative summary effect sizes at each level of sampling variation (observation error).

The summary effect size, and summary interactions class as generated from the meta-analytical framework also shows some sensitivity to observation error, although in these analyses the outcome is rather different (Figure 2b). For low levels of observation error, the 95% confidence intervals of the summary effect size overlap zero, indicative of an additive summary interaction class. This occurs because the frequency and magnitudes of synergistic (positive effect size) and antagonistic/reversal (negative effect sizes) interactions are approximately equal for low observation error (Figure 2a), and although there is a large variance in effect sizes due to low sampling error (See Supplementary Material 1), the effects sizes for individual interactions are approximately centred on zero. However, with increasing observation error the summary effect sizes become increasingly more negative, and confidence intervals for these summary effect sizes do not overlap zero, indicating an antagonistic/reversal summary interaction class. Further inspection shows an increase in the proportion of negative effect sizes as observation error increases (Figure 2c), with this being mirrored by a decreasing summary effect size (Figure 2b). Although not so obvious due to the dominance of additive interactions, a similar trend can be observed in the frequencies of interaction types at higher observation errors, with synergistic interactions heading towards 0 frequency faster than antagonistic interactions (Figure 2a). Hence, analyses of our model results with varying levels of observation error suggest synergies in pairs of ecosystem stressors may be under-reported in many empirical studies.

### Theoretical predictions

In summary, our theoretical analyses lead us to predict that at likely levels of sampling variation we should expect the empirical data to be dominated by additive interactions for individual interactions (Figure 2a), but in contrast the summary effect sizes computed across a large body of such studies should indicate a dominant role for antagonistic, or reversal, interactions.

### Stressor Interactions within Freshwater Empirical Data

Our literature search within Web of Science returned 1805 papers that meet our search criteria. Of these, only 58 meet our criteria for inclusion. They include 546 interactions summarised in Figure 3 to show the frequency of different interaction classifications and the overall summary effect sizes and interaction classes. Additive interactions were the most frequent, (0.830), followed by antagonistic, (0.0989), reversal, (0.0476), and finally synergistic, (0.0238), interactions (Figure 3a).

**Figure 3:**
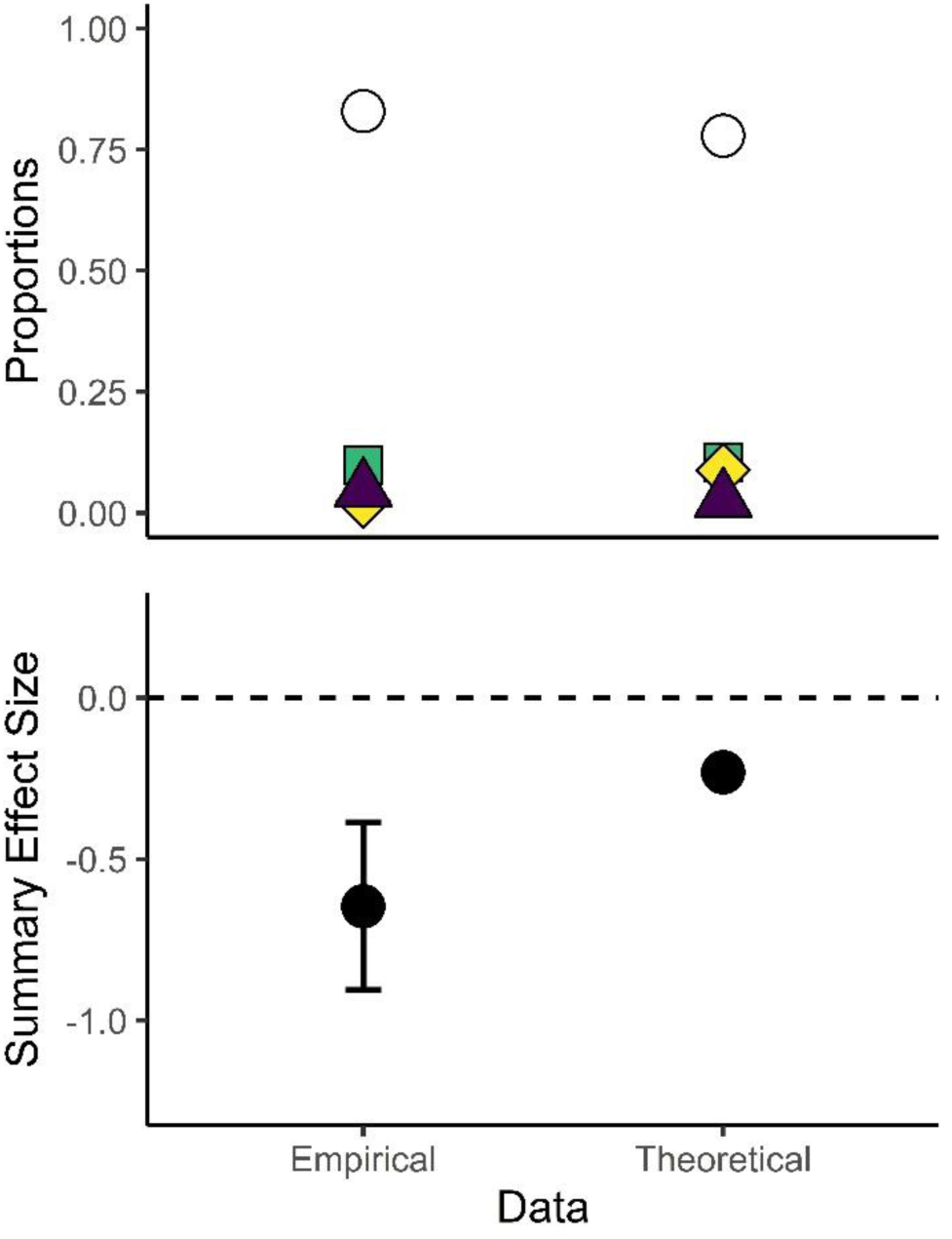
a) Proportions of the different interaction classes, and b) summary effect sizes for the empirical and theoretical dataset. The empirical dataset comprised 546 interactions, while the theoretical dataset comprised all interactions using sampling variations between 1×10^−2^ and 0.5. (5,040,000 interactions). White circles denote additive interactions. Green squares denote antagonistic interactions. Yellow diamonds denote synergistic interactions. Purple triangles denote reversal interactions.

Additionally, the summary effect size for the entire dataset is negative, (−0.646 ± 0.259), with confidence intervals that do not overlap zero, indicative of an antagonistic/reversal summary interaction class (Figure 3b).

Our meta-analysis reports medium-level heterogeneity, (I^2^ = 47.0%), though this is considerably lower than the mean heterogeneity, (I^2^ = 91.7%) found in an analysis of previous ecological meta-analyses (Senior et al., 2016). Two additional meta-analyses, conducted upon sub-groups of the empirical dataset, with the categorical moderator of organism group as means of exploring this heterogeneity (Nakagawa et al., 2017) fail to uncover any source of this heterogeneity (see Supplementary Material 6).

### Comparison of Empirical and Theoretical Interaction Classifications

Overall, we find close agreement between our theoretical models with biologically reasonable levels of observation error and the freshwater empirical data (Figure 3). Summary effect sizes are negative indicating antagonistic or reversal interactions (Figure 3b); whereas the vote counting results highlight how individual interactions tend to return an additive classification (Figure 3a), probably due to the sampling errors, and relatively low sample sizes, in both data sets.

## Discussion

There has been much interest in understanding and cataloguing the joint effects of stressors on ecological communities and ecosystems (Schäfer & Piggott 2018; Thompson et al. 2018b), but to date there has been relatively little guidance from ecological theory. Here we close this gap by analysing a food chain model using classical ecological theory and comparing it to a meta-analysis on a large dataset for freshwater ecosystems. Our theoretical results show remarkable agreement with the empirical analyses, for both vote counting results and summary effect sizes, which generate different interpretations of how stressors are likely to interact (Figures 2, 3). On the one hand, our vote counting analyses suggest additive interactions to be by far the most dominant stressor interactions in freshwater communities; but on the other hand our meta-analysis shows antagonism to be the summary interaction class. Our theoretical model helps to understand why this might be the case, and highlights deficiencies in the commonly used additive null model that is used to classify the joint effects of ecosystem stressors. In particular our model results show the additive null model is (a) sensitive to sampling variation with even realistically small levels leading to very frequent failure to correctly reject the null model (type II statistical errors); (b) potentially less likely to correctly report synergistic interactions compared to either antagonistic or reversal interactions in the meta-analytical framework. We believe that once these statistical aspects are considered, so-called ‘ecological surprises’ (*sensu* Paine et al. 1998) may in fact be more prevalent in both our freshwater dataset, and more widely.

### Theoretical predictions

The agreement between theoretical models and empirical data is remarkable given the biological simplicity of the model and how it is not tailored to any one type of stressor or community. However, our approach should be viewed as one that aims to explain the emergent patterns across studies rather than be used to predict the joint effects of stressors in a particular empirical system, in which case a more detailed and specific model is more appropriate (e.g. Brown et al. 2013; Galic et al. 2018). Our food chain models imply that, given adequate sample sizes (see below), we should expect synergistic and antagonistic interactions to co-dominate at the population and trophic level, whereas additive interactions and reversals should be relatively rare. These messages appear to be echoed in the few other theoretical studies on stressor interactions in ecological communities (e.g. Travers-Trolet et al. 2014; Thompson et al. 2018a; Haller-Bull & Bode 2019). This agreement is despite a variety of key differences in the model assumptions. In particular, Haller-Bull and Bode (2019) focussed on populations rather than multispecies communities, but found dominant roles for synergistic and antagonistic interactions, with additive interactions occurring most frequently for stressors affecting the carrying capacity. Similar to our model, Thompson et al. (2018a) did focus on multispecies communities, but they assumed biological interactions were constant, whereas we allow interactions (consumption and conversion rates) to be modified by stressors, an assumption that seems likely to be met on a regular basis. For example, stressors have been shown to influence resource competition (Kroeker et al. 2013); susceptibility to parasitism in oysters (Lenihan et al. 1999); and modify the flow of energy through aquatic food webs by inducing changes in trophic links (Schrama et al. 2017). Despite this difference, Thompson et al. (2018a) found additive interactions were most prominent when species facilitated one another (i.e. positive species interactions), but that synergy or antagonism in combined stressor effects on species richness or community biomass were more common when species interactions are negative (competition or resource use). Finally, Fu et al. (2018) used four ecosystem models for fisheries to investigate the combined effects of fishing and primary productivity across a number of modelled real-life fisheries. They also found a reduced role for additive interactions, with an increased risk of stressor pair synergism at lower trophic levels, whereas antagonistic interactions (less than additive, but in the same direction as the additive expectation) where more likely at higher trophic levels.

The apparent rarity of additive interactions in all of these models might appear at odds with the possible interpretation that two stressors acting on different species within a community could lead to such a joint effect (Jackson et al. 2016). However, feedbacks in the food web, like those found in our models, mean that even if a species is unaffected directly by a stressor, it is highly likely that top-down or bottom up effects will lead to indirect interactions for many species, and as a result additive interactions are extremely hard to generate in the absence of sampling variation (e.g. observation error). Indeed, we predict that additive interactions might only truly occur in scenarios where species in different and very weakly interacting sub-communities are affected by different stressors, or, as found by Thompson et al. (2018a), where species interactions are predominantly positive. However, despite a growing body of theoretical predictions we are not aware of any empirical test of the previous models. Our models have therefore extended earlier results by focussing on changes in biological interactions caused by the stressors, and also incorporating sampling variation as a parameter of interest, something that greatly aided the interpretation of the empirical results. We believe there will be an increasing role of theory in generating hypotheses for the ways in which stressors interact (De Laender 2018), and the most progress will be made when the theory is developed so it can be tested directly against the data, much as we have done here.

### Sample size

The choice of null model is hotly debated within ecological stressor research (Schäfer & Piggott 2018), and it has been argued that null models should be able to accurately predict the combined effects of stressors (Orr et al. 2020). However, our work does add some cautionary notes to this view since it is clear that the additive null model for stressor interactions is very sensitive to sampling variation, and for likely realistic levels of sampling variation it is hard to correctly reject the null model (Figure 2). Given that most experiments have low sample sizes (a mean of 3.83 with a maximum of 16 per treatment in our empirical data), we feel it is premature to conclude that most stressor interactions are *truly* additive in the freshwater data we collected. This view is reinforced by our meta-analysis that returned a negative summary effect size implying an overall antagonistic, or reversal, summary interaction class within in experimental results, a pattern that was mirrored in previous analyses of freshwater stressor experiments (Jackson et al. 2016; Lange et al. 2018). Also, given that our theory, in the absence of sampling variation, showed a near equal frequency of synergistic and antagonistic interactions (Figure 2a), there appears to be a potential trend against detecting synergies in co-occurring stressors in the meta-analytical framework (Figure 2b). The dual effects of this potential trend and sensitivity to sampling variation may be key reasons why stressor synergies are not as often reported as might be expected (Darling and Côté 2008; Côté et al. 2016), although as we discuss below, other reasons may also contribute, and of course, we cannot rule out that the empirical results do truly reflect the underlying interactions. However, our finding of sensitivity to sample size is more general than either our theoretical results, or our freshwater dataset, and we suggest future work should investigate other null models for their robustness to these (and other) features. For example, is the additive null model particularly conservative in its detection of synergies, and are there better alternatives? Such analyses would build on previous descriptions of the null models (e.g. Sih et al. 1998; Folt et al. 1999; Sih et al. 2004) and would be particularly useful if analyses considered the effect of sample size on statistical power, as this will help guide future empirical studies to improve the detection rate of non-null stressor interactions. Furthermore, a previous theoretical analysis, implementing an alternate framework to that used here, found that synergistic interactions only occurred under specific conditions (Haller-Bull & Bode 2019). Accordingly, future theoretical studies may wish to investigate the controls that govern the frequency of synergistic interactions, and in doing so determine whether such patterns are general or more tailored to specific models. Overall, it is important to note that when comparing observed interactions to a null model we are determining whether it is possible to reject the null model. Similarly, a failure to reject the null model does not mean that the stressors interact in an additive manner, only that we are unable to find a statistically significant difference between what is observed and what is predicted. Ultimately, acknowledging the difference between these two statements, and the corresponding interpretation of a null model, is crucial when attempting to further our collective understanding of these statistical tools.

### Lack of generalities across meta-analyses

Very few general patterns have emerged from previous meta-analyses on stressor interactions (Côté et al. 2016), but there are a number of reasons as to why this is the case (see also Côté et al. 2016). Firstly, the studies have been carried out across all the different major realms (marine, terrestrial, and freshwater) and there could be heterogeneity simply because different stressor interactions might prevail in the different realms. Secondly, there is both a range of stressors considered, and a naturally large taxonomic variation in study organisms cutting across a wide range of life histories and trophic structures. For example, it could be that long-lived and short-lived organisms experience different effects, or for instance, that trophic level is important to the type of stressor interaction that tends to occur (Thompson et al. 2018a; see Supplementary Material 5), and that different combinations of stressors will give rise to different forms of interaction (e.g. Jackson et al. 2016, see Supplementary Material 5). Thirdly, different meta-analyses have considered different levels of biological organisation, from individuals, to populations communities and ecosystems (reviewed by Crain et al. 2008 for marine ecosystems), and we can expect different interactions to occur for the same stressor pair across the levels of organisation (e.g. Galic et al. 2018). Fourthly, there is a profusion of null models and classification schemes for stressor interactions (Schäfer & Piggott 2018; Orr et al. 2020), making comparisons between studies very difficult, especially when we do not know the relationships between different null models. For example, under the same dataset, when should we expect synergistic and antagonistic interactions to be reclassified when we move from, say, the additive null model, to the multiplicative null model? Finally, we note that there is variation in the statistical methodologies implemented across meta-analyses. For instance, the manner in which interactions are classified can vary between methodologies (e.g. Crain et al. 2008 versus Darling & Côté 2008) which may potentially result in contrasting frequencies of the different interaction classifications being reported. We believe the first step to uncovering any generalities across meta-analyses is to eliminate any roles that methodological differences are playing, and only then can we focus on the more interesting biological causes (i.e. sources 1-4) for similarities and differences in the ways multiple stressors combine across different ecological communities.

### Mechanistic understanding of multiple stressors

Here, we sought an to answer to the question of *how* multiple stressors interact. This approach, when applied across both theoretical and empirical datasets can allow us to discern what might be expected across the interactions of multiple stressors. However, future research may seek to answer the question of *why* multiple stressors interact in the manner that they do. Undoubtedly, these two questions are entwinned, with the answers to each of these questions highly likely to be dependent upon the other. However, while the use of null models is essential in determining the combined effect of multiple stressors (Thompson et al. 2018b), the adoption of a mechanistic approach to investigating multiple stressors may provide novel insights which address these joint questions (De Laender 2018; Schäfer & Piggott 2018). For instance, a mechanistic understanding may allow for responses such as co-tolerance or co-susceptibility (Todgham & Stillman 2013) to stressors to be more thoroughly understood from an ecological perspective. Ultimately, such an understanding is likely to require a large amount of empirical data to fully understand; however, there is ample scope for theoretical ecology to help fill this gap in our collective understanding of multiple stressors, and to generate specific hypotheses to be tested. Similarly, a mechanistic understanding of multiple stressor interactions would prove invaluable when mitigating the effects of stressors or implementing conservation initiatives.

### Conclusions

Here we have detailed the first empirical test of general theoretical predictions for how multiple stressors interact across a large number of freshwater community case studies. Our empirical results suggest that additive interactions are pervasive at the study level, but that meta-analyses reveal a summary antagonistic, or reversal, interaction class for the entire freshwater community dataset. However, our theory suggests these results may be reflecting sampling variation rather than any underlying stressor interaction, and that so-called ecological surprises may be far more common than empirical analyses are suggesting, with the theoretical results indicating similar frequencies of antagonistic and synergistic interactions. Predicting the ways multiple stressors interact is key when attempting to mitigate their effects, with the class of observed interaction potentially outlining whether the removal of a stressor will have a beneficial, limited, or detrimental impact to the system (Brown et al. 2013; Côté et al. 2016). Our results show the value of developing a theoretical framework for predicting and understanding environmental stressor interactions, and we hope more general theory that makes specific predictions based upon ecological mechanisms (e.g. De Laender 2018; Fu et al. 2018; Thompson et al. 2018a) will be developed *and* tested in the future. However, our results also highlight the need to better understand the strengths and limitations of the null models that are used to test classify the cumulative effects of community stressors, and we also believe a unified approach to the meta-analyses of individual studies will increase our understanding of how environmental stressors combine.

## Acknowledgements

We thank Rory Gibb, Michelle Jackson, and Tim Newbold for thought-provoking discussions. This research was part-funded by the Natural Environment Research Council grant NE/M010481/1.

