## Supplementary Material for "Ecological theory predicts ecosystem stressor interactions in freshwater ecosystems, but highlights the strengths and weaknesses of the additive null model"

**Supplementary Material 1** – *Theoretical Simulations*

The following section details the methodology used to generate data for the theoretical simulations.

*Equations*

The equations, (Equations 1 & 2), used throughout this analysis have been taken from Heath et al., (2014); though the equations have been minorly adapted to include density independent mortality at all trophic levels, (except for the bottom trophic level), as opposed to only the apex trophic level. The two sets of equations comprise differing forms of dynamics and regulation. Equation 1, (DD-Equations), comprises a term for density-dependent mortality; whereas in Equation 2, (CR-Equations), consumer uptake is regulated by density. These two sets of equations have been explored in detail, and compared to alternate forms of classic ecological theory, by Heath et al. (2014).

*Generation of Theoretical Data*

Theoretical stressor interaction data was generated using the following methodology, (expanded from that in the Methods section).

1) Parameter combinations were randomly determined for 1,320,000 food chains; where each combination of equation type, (DD-Equations or CR-Equations), and food chain length, (three, four, or five trophic levels), accounts for 220,000 parameter combinations. Each parameter was randomly assigned a value from a predetermined range, with ranges differing depending upon the parameter in question, (Table 2). The parameter values were determined using a uniform distribution, i.e. any value in the range is as likely to be selected as any other. These values are considered to be *baseline* parameter values.

2) Two parameters were randomly chosen to be subjected to stress with all candidate parameters being chosen with equal probability. These two parameters have different values to their baseline

equivalents, but all other parameters will maintain the same values. Stressed parameter values were chosen from the ranges given in Table 2, using a uniform distribution. Only the parameters of attack rate (α), conversion efficiency (ε), density independent mortality (δ), or nutrient input rate (ω) were able to be stressed. The parameters selected to be stressed could be on the same trophic level, (e.g. $\alpha_2$ and $\varepsilon_2$), the same biological process on different trophic levels, (e.g. $\varepsilon_2$ and $\varepsilon_1$), or different biological processes on different trophic levels, (e.g. $\varepsilon_1$ and $\alpha_2$). Additionally, $\alpha_1$ was not stressed in any of the simulations. This is due to the fact that altering $\alpha_1$ only had any impact upon the nutrient trophic level, and no impact upon any other trophic levels in the simulations, (unpublished data). Given that this analysis was primarily concerned about *non-nutrient trophic levels*, $\alpha_1$ was not stressed in any analysis.

*Stressor A* parameter values are the same as the baseline values except that the parameter value for the first stressed parameter is changed to the stressed value.

*Stressor B* parameter values are the same as the baseline values except that the parameter value for the second stressed parameter is changed to the stressed value.

*Interaction* parameter values are the same as the baseline values except that the parameter value for both stressed parameters are changed to the corresponding stressed values.

Overall, there are 5,280,000 parameter combinations equating to 1,320,000 parameter groups for individual food chains under baseline, stressor A, stressor B, and interaction conditions.

3) For each of the 1,320,000 parameter groups, the equilibrium densities of all trophic levels in the food chain, under all levels of stress, were determined using the *NSolve* function in Mathematica 10.4, (Wolfram Research Inc., 2016). Equilibrium densities are referred to as the densities of each trophic level in the food chain that result in there being no net change in the density of any other trophic level. Furthermore, additional constraints were placed on the equilibrium densities in that

they needed to be feasible, (i.e. the equilibrium densities of all trophic levels in the food chain had to be positive and non-zero).

For some parameter groups, equilibrium densities were able to be calculated for the baseline parameter values but not for either stressor A, stressor B, or the interaction parameter values. When this occurred the equilibrium densities for the entire parameter group were discarded from any further analysis.

Overall, feasible equilibrium densities could be found, under all four levels of stress, for 1,054,272, (79.9%), parameter groups.

4) For each of the 1,054,272 parameter groups the stability of the food chains was assessed. This was concluded by taking the second differential of Equations 1 and 2. Overall, in order for the equilibrium densities of each food chain, (under each level of stress), to be considered stable, the following conditions had to be met, outlined in Equations S1 and S2. Of the 1,054,272 parameter groups, all food chains, under all levels of stress, were stable.

$$\frac{d^2x_0}{dt^2} : 0 > -\alpha_1 x_1$$

$$\frac{d^2x_i}{dt^2} : \alpha_{i+1}x_{i+1} + \delta_i + 2\,\lambda\,{}_i x_i > \alpha_i \varepsilon_i x_{i-1}$$

$$\frac{d^2x_n}{dt^2} : \delta_n + 2\,\lambda\,{}_n x_n > a_n \varepsilon_n x_{n-1}$$

Equation S1: Conditions that need to be met in order for each trophic level in a community, (with DD-Equations), to be considered stable. Notation is the same as in Table 1.

$$\frac{d^2x_0}{dt^2} : 0 > -\frac{\alpha_1 x_1}{(1 + v_1 x_1)}$$

$$\frac{d^2x_i}{dt^2} : \frac{\alpha_{i+1}x_{i+1}}{(1 + v_i x_i)} + \delta_i > \frac{(\alpha_i \varepsilon_i x_{i-1})(1 + v_i x_i) - (v_i)(\alpha_i \varepsilon_i x_{i-1} x_i)}{(1 + v_i x_i)^2}$$

$$\frac{d^2x_n}{dt^2}:\ \delta_n > \frac{(\alpha_n\varepsilon_n x_{n-1})(1+\ v_n x_n) - (v_n)(\alpha_n\varepsilon_n x_{n-1}x_n)}{(1+\ v_n x_n)^2}$$

Equation S2: Conditions that need to be met in order for each trophic level in a community, (with CR-Equations), to be considered stable. Notation is the same as in Table 1.

5) From the 1,054,272 parameter groups, 360,000 were randomly selected, (with each combination of food chain length and equations accounting for ~16.667% of interactions). From each of these 360,000 parameter groups, the equilibrium density of a single trophic level, (consistent across each level of stress), was randomly selected and collated into a single dataset for analysis, hence referred to as the *analysis dataset*. All subsequent analysis was performed on this analysis dataset.

*Simulation of Observation Error*

For each parameter group, when calculating the equilibrium densities, there was no stochasticity included within the equations. Hence, using the same parameter values will return the same equilibrium densities each time. However, this is not necessarily reflective of ecological systems. To account for this, stochasticity, (through the form of observation error), was incorporated into the analysis dataset through the following methodology.

6) Firstly, the level of observation error to be included was chosen. Within this analysis, the level of observation error refers to the standard deviation of a normal distribution with a mean of 1. Within this analysis, observation was included with equal spacing on a logarithmic scale, (between $1\text{x}10^{-10}$ and $5\text{x}10^{-1}$), with there being 86 different levels of observation error investigated, (alongside the absence of observation error). Through the rest of this methodology we use a level of observation error of σ.

7) For each of the 360,000 parameter groups in the analysis, a number of replicates, (*n*), (either three, four, five, or six), was randomly selected using a uniform distribution. The number of replicates was consistent across the different treatments for each parameter group, (i.e. baseline, stressor A, stressor B, and interaction). The rationale for choosing a number of treatments between three and six was that these are sample sizes frequently seen in empirical studies, and as such making this theoretical analysis akin to empirical studies in this regard.

8) For each treatment within each parameter group, *n* values from a Gaussian distribution with a mean of 1 and standard deviation of σ were taken. These values were then multiplied by the equilibrium density for that treatment; with the mean and standard deviation of these new *n* equilibrium densities being calculated.

   This was conducted for each treatment within each parameter group. Additionally, if any of the *n* values from the Gaussian distribution were equal to, or greater than, 2 or less than, or equal, to 0, then the *n* values from the Gaussian distribution were re-drawn.

   Accordingly, by having the mean, standard deviation, and number of replicates for each treatment, it is possible to calculate effect size metrics, (Hedges' d), and determine interaction classifications, (see Supplementary Material 3).

*Additional Results*

Overall, the qualitative pattern for the interaction classification frequencies varied with observation error. Figure S1.1 shows how the frequency of the different interaction classes varies with observation error for the overall dataset of 360,000 interactions, as well as this dataset when subset for food chains using either CR-Equations or DD-Equations. Similarly, Figure S1.2 shows how the frequency of interaction classes varies when the overall dataset is subset by the length of food chain. Additionally, Figure S1.3 shows how interactions classed as being reversal, antagonistic, or synergistic under no observation error, are reclassified when observation error is incorporated at different levels.

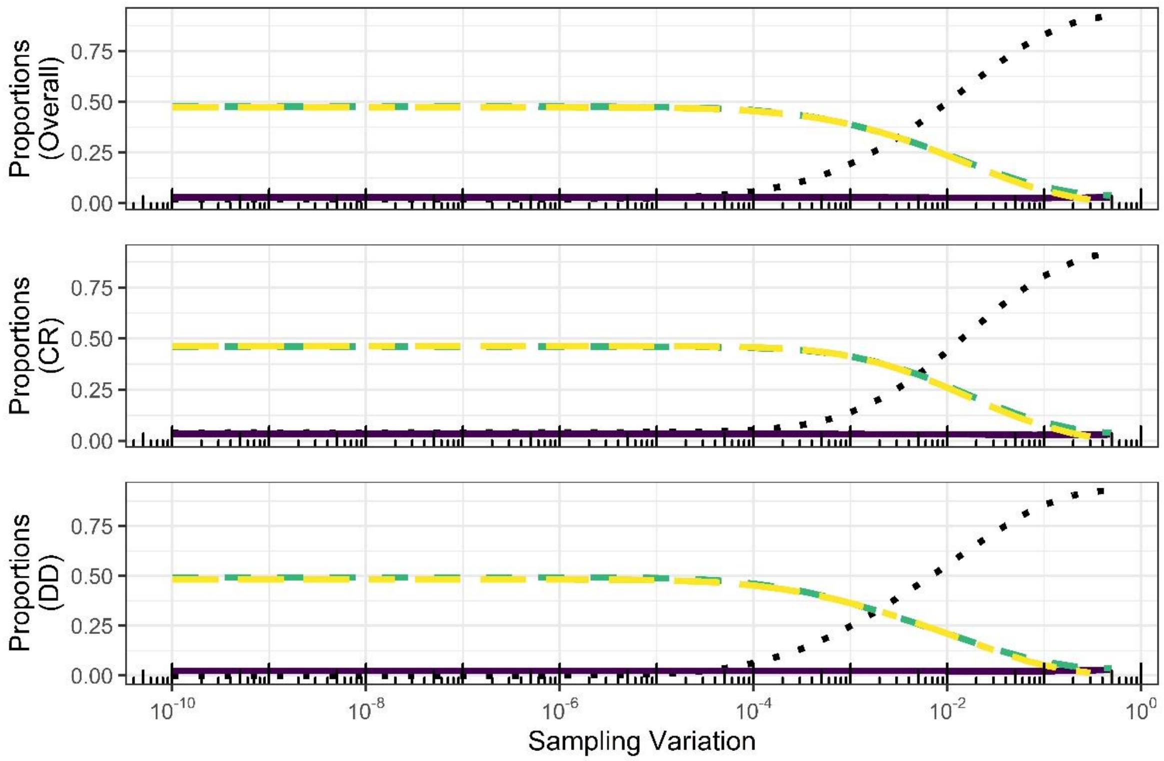


Figure S1.1: Prevalence of the different classification of interaction for the theoretical data, when observational error was implemented, separated for data from Consumer Uptake Regulation Equations, Density Dependence Equations, and the combined dataset. Observation standard deviation denotes the standard deviation that was used to determine observation error. Top – Overall dataset, Middle – only food chains with CR dynamics, Bottom – only food chains with DD dynamics. Dotted black line denotes additive interactions. Dashed green line indicates antagonistic interactions. Long-dashed yellow line denotes synergistic interactions. Solid purple line indicates reversal interactions.

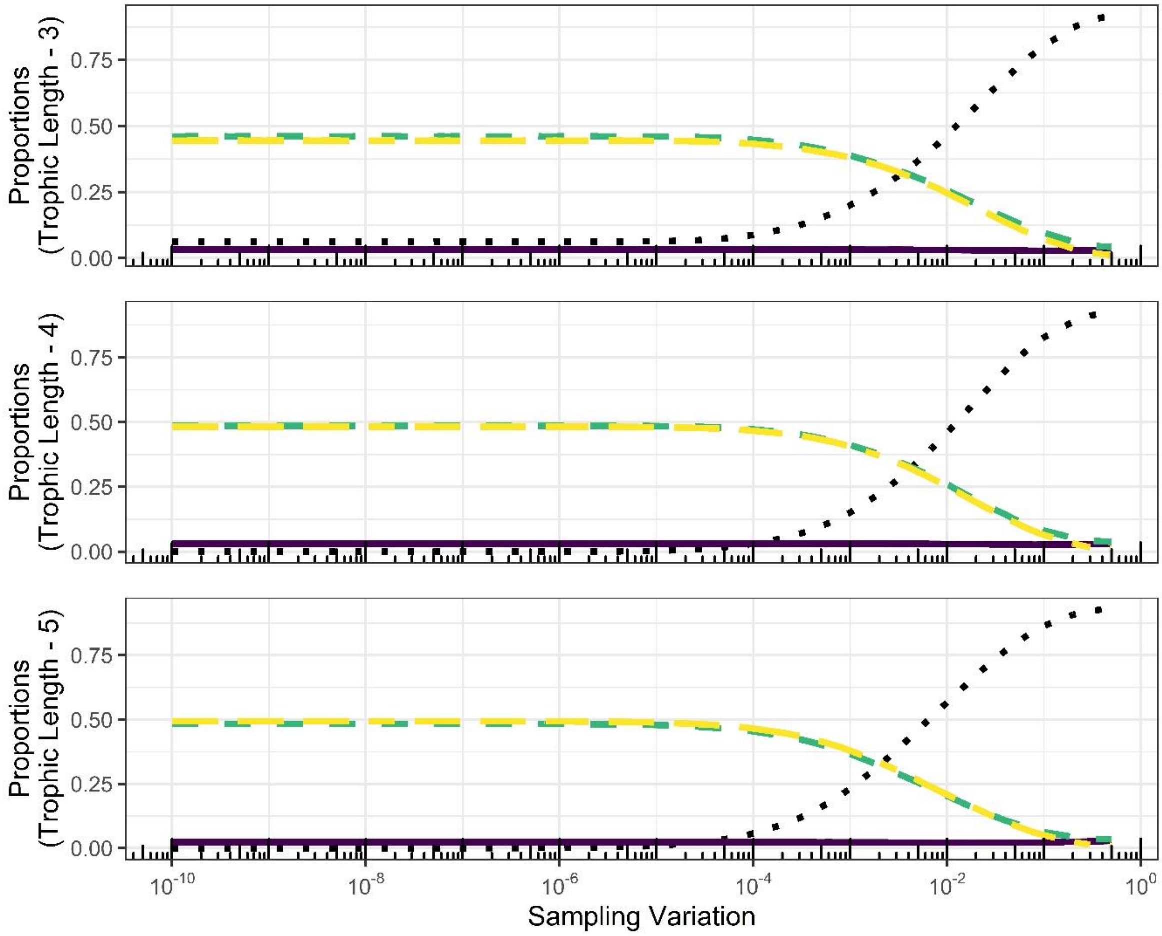


Figure S1.2: Prevalence of the different classification of interaction for the theoretical data, when observational error was implemented, separated for data from Consumer Uptake Regulation Equations (Equations 1, main text), Density Dependence Equations (Equations 2, main text), and the combined dataset. Standard deviation denotes that which was used to determine observation error. Top – Communities comprising three trophic levels, Middle – Communities comprising four trophic levels 4, Bottom – Communities comprising five trophic levels. Dotted black line denotes additive interactions. Dashed green line indicates antagonistic interactions. Long-dashed yellow line denotes synergistic interactions. Solid purple line indicates reversal interactions.

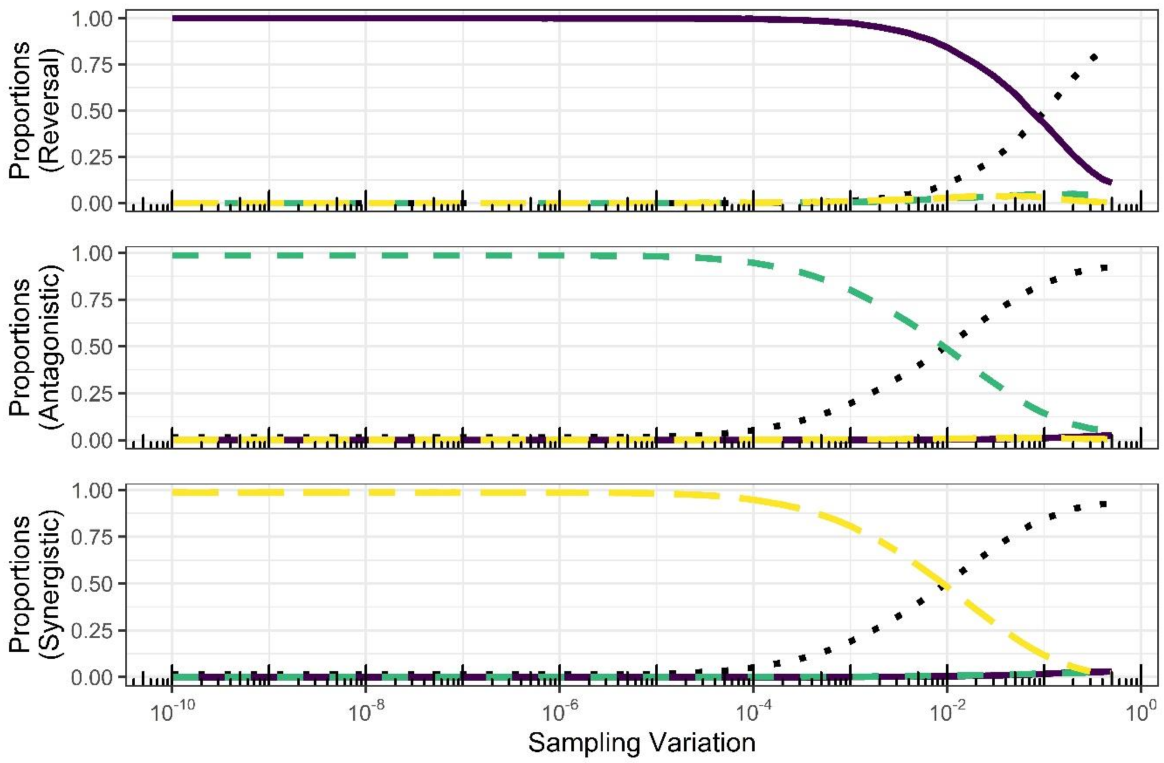


Figure S1.3: For a given 'true' interaction classification, (i.e. when observation error is not considered), the proportions of this interaction classification that are reassigned to a different class when observation error is implemented. The data shown is for the combined dataset of containing simulations using either set of equations, comprising 360,000 interactions. Top – Reversal interactions, Middle – Antagonistic interactions, Bottom – Synergistic Interactions. Dotted black line denotes additive interactions. Dashed green line indicates antagonistic interactions. Long-dashed yellow line denotes synergistic interactions. Solid purple line indicates reversal interactions.

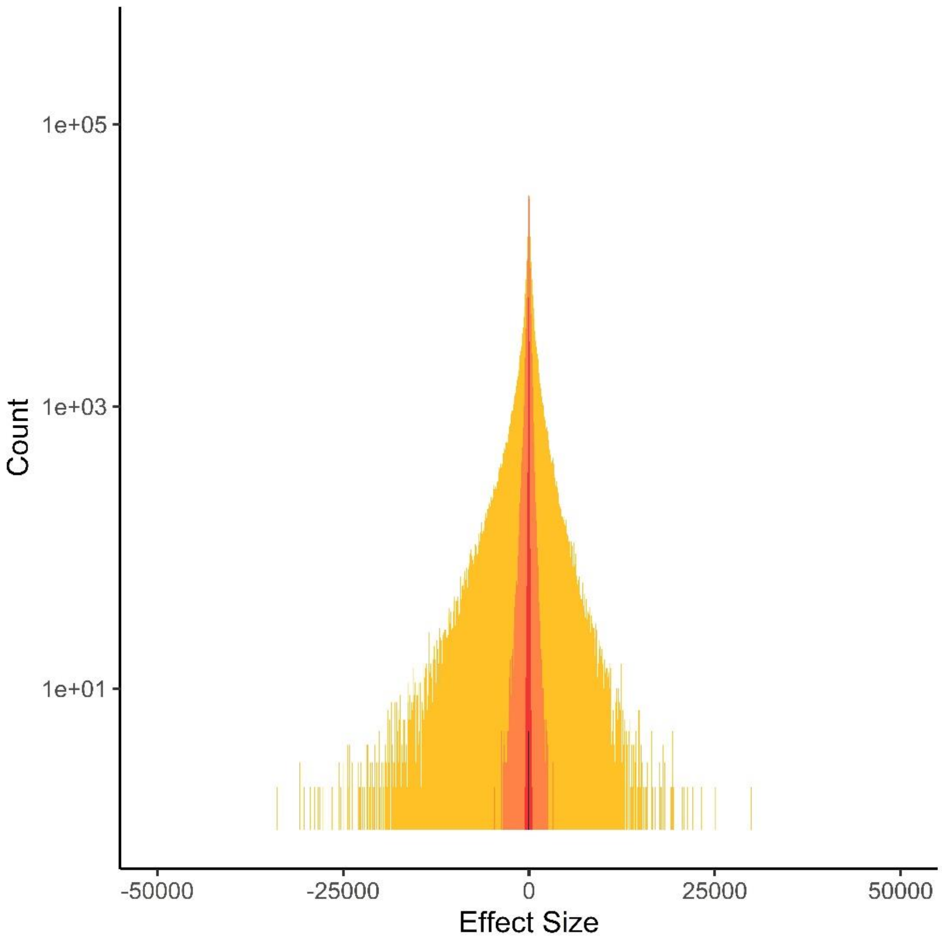


Figure S1.4: Distribution of effect sizes for the theoretical data, under multiple levels of sampling variation. Yellow indicates the effect size distribution for a sampling variation of $1x10^{-4}$; Orange indicates the effect size distribution for a sampling variation of $1x10^{-3}$; Red indicates the effect size distribution for a sampling variation of $1x10^{-2}$; Black indicates the effect size distribution for a sampling variation of $1x10^{-1}$.

Wolfram Research. Inc., 2016. Mathematica, (Version 10.4). *Wolfram Research, Inc., Champaign, Illinois.*

**Supplementary Material 2** – *Literature Search*

A search for literature was conducted in Web of Science, (26/02/2019), using the following search criteria, for papers published up to, and including, 31st December 2018:

TS = ("synerg*" OR "antagon*" OR "additive" OR "dampen*" OR "combined" OR "multiple" OR "factorial" OR "experiment" OR "stressor" OR "multiple stressor") AND TS = ("freshwater" OR "river" OR "stream" OR "lake" OR "catchment" OR "pond" OR "riparian") AND TS = ("impact" OR "effect") AND TS = ("communit*") AND TS = ("biomas*" OR "abundan*" OR "densit*") AND SU=("Biodiversity & Conservation" OR "Environmental Sciences & Ecology" OR "Fisheries" OR "Marine & Freshwater Biology" OR "Mathematical & Computational Biology" OR "Microbiology" OR "Plant Sciences" OR "Zoology")

The search terms used here were adapted from those used by Jackson et al. (2016).

Overall 1805 entries were returned.

To be included within this dataset, studies had to investigate the impacts of two stressors, upon a freshwater community. There was no limit placed upon the types of stressor that could be considered, so long as the stressor was a manipulation of environmental, or novel ecological, conditions as opposed to measuring the response of native communities to normal, or expected, events or disturbances.

Only data referring to the abundance, biomass, density, (or in some cases chlorophyll-a, or other chemical concentrations from which biomass could be estimated), was collated.

For each group of organisms, the study had to report, mean values, standard deviation or standard error, and the number of replicates for the following:

i) Data for the group of organisms under control conditions, (i.e. no stressors)

ii) Data for the group of organisms under Stressor A, (a single stressor)

iii) Data for the group of organisms under Stressor B, (a single stressor)

iv) Data for the group of organisms under both Stressors A and B, (both stressors acting simultaneously)

Each study could report data for multiple different groups of organisms, all of which was collated. Additionally, some studies reported data for freshwater communities exposed to multiple different intensities of a stressor, all of these were collated, so long as the above factorial design was followed. Where studies did not report mean values, standard deviation or standard error, or the number of replicates, the data was not incorporated within the dataset.

Each paper was individually screened to determine whether it was suitable for inclusion within our study. Firstly, titles and abstracts were assessed to determine whether the paper was potentially relevant to our study. If a paper was potentially relevant the methods were read to determine whether the study design was factorial and met the above criteria for inclusion. If the paper met the criteria, then the results and supplementary material were considered to determine whether the all required data, (above), was presented. Overall, data extraction was attempted for 124 papers. However, at this stage 66 papers were deemed unsuitable for data extraction, due to issues such as missing data, or having figures that were too unclear for data extraction.

To be included within this dataset, studies had to investigate freshwater communities. Those were experiments were conducted upon other such aquatic environments, e.g. marsh, brackish, estuarine, or coastal environments were not considered.

From each study, data was extracted from tables or from figures using WebPlotDigitizer software, (Rohatgi 2017).

Overall, 58 papers were found, which were able to contribute data to this dataset, contributing a total of 859 interactions. However, to reduce covariance within the dataset between these interactions, the following protocol was adopted.

i) Where studies reported data for multiple response metrics, preference was given in the following order: Density, Abundance, Biomass, and Chlorophyll-a (and other chemical proxies for biomass).

ii) Where studies reported data for multiple overlapping groups of organisms, preference was given to subsets of groups of organisms as opposed to coarser groupings. For instance, if a study reported densities of i) overall periphyton, alongside data for constituent ii) algae and iii) bacteria; only ii and iii would be included in the dataset.

iii) Where studies reported data for the same group of organism, but in different locations within the study environment, only one location was used, with preference being given to the location were a more complete set of data was able to be extracted.

iv) Where studies reported data for multiple time points, preference was given to the time point measured latest in the experiment.

After following this protocol, the number of interactions was reduced to 546.

Within the dataset, each stressor is assigned into one of seven different categories depending upon qualitative properties of the stressor. The seven different categories are: Community Composition, Contamination, Habitat Alteration, Light, Nutrients, Salinity, and Temperature, with explanations and examples given in Table S2.

Table S2: The seven broad categories of stressor considered within this analysis, with examples and definitions provided.

| Stressor Category | Explanation | Example |
|---|---|---|
| Community Composition | The addition or removal of a group of organisms from the community. | e.g. Invasive species |
| Contamination | The addition of a chemical, (with the exception of nutrient), at concentrations not normally found in the system. | e.g. Pesticides |
| Habitat Alteration | Changing the physical properties of a system from those normally seen | e.g. Sedimentation |
| Light | Change in the levels of light that a system receives | e.g. UV |
| Nutrients | Addition of nutrients to a system at levels exceeding those normally found | e.g. Nitrogen |
| Salinity | An increase in the salinity of the freshwater study system | e.g. Salt addition |
| Temperature | An increase in the temperature of the freshwater study system | e.g. Temperature increase |

Overall, Hedges' d, ($d_{int}$), was calculated using the following equations:

$$d_{int} = \frac{D_I - D_A - D_B + D_C}{s}$$

Where $s$ is the pooled standard deviation of $d_{int}$:

$$s = \sqrt{\frac{(N_I - 1)(s_I)^2 + (N_A - 1)(s_A)^2 + (N_B - 1)(s_B)^2 + (N_C - 1)(s_C)^2}{(N_I + N_A + N_B + N_C - 4)}}$$

The variance of $d_{int}$ can be calculated as:

$$v_{int} = \left(\frac{1}{N_I} + \frac{1}{N_A} + \frac{1}{N_B} + \frac{1}{N_C} + \frac{d_{int}^2}{2(N_I + N_A + N_B + N_C)}\right)$$

Where the standard error of $d_{int}$ can be calculated (Borenstein 2009) as:

$$SE_{int} = \sqrt{v_{int}}$$

Where 95% confidence intervals are calculated as:

$$CI_{95\%} = 1.96 * SE_{int}$$

For each interaction, directionality was removed using the methodology stated in the Methods section, outlined in Piggott et al. (2015). This involved inverting the response direction of the calculated Hedges' d where the predictive additive effects were negative.

For each interaction, it is possible to use Hedges' d to determine an interaction classification, (namely, additive, antagonistic, synergistic, or reversal). Accordingly, an interaction was classified as being additive if, $d_{int} \pm CI_{95\%}$, overlaps zero. If $d_{int} \pm CI_{95\%}$ does not overlap zero, then an interaction was classified under the following conditions:

i) $d_{int} > 0$, the interaction is classed as being synergistic

ii) $d_{int} < 0$ and

$D_A + D_B - 2D_C$ and $D_I - D_C$ have the same polarity, the interaction is classed as being antagonistic

iii) $d_{int} < 0$ and

$D_A + D_B - 2D_C$ and $D_I - D_C$have the different polarities, the interaction is classed as being reversal

The methodology was used for all empirical interactions, and for theoretical interactions where observation error was included.

*Theoretical Interactions without observation error*

When no observation error was present in the theoretical experiments, $d_{int}$ was instead classified using the following equation:

$$d_{int} = D_I - D_A - D_B + D_C$$

This is due to the fact that with no observation error, the was no variation in the densities of trophic levels, hence effect sizes could be determined exactly with no confidence intervals.

Accordingly, under this set of scenarios, if:

iv) $d_{int} > 0$, the interaction is classed as being synergistic

v) $d_{int} < 0$ and

$D_A + D_B - 2D_C$ and $D_I - D_C$ have the same polarity, the interaction is classed as being antagonistic

vi) $d_{int} < 0$ and

$D_A + D_B - 2D_C$ and $D_I - D_C$ have the different polarities, the interaction is classed as being reversal

vii) $d_{int} = 0$, the interaction is classed as being additive

Table S4: Comparisons of the naming of interaction classifications within this study, and other studies.

| This research | Travers-Trolet et al. (2014) / Fu et al. (2018) | Common Nomenclature |
|---|---|---|
| Additive | Additive | Additive |
| Synergistic | Synergistic | Synergistic |
| *Antagonistic* | *Dampened* | *Antagonistic* |
| *Reversal* | *Antagonistic* | *Antagonistic* |

The term *antagonistic* is frequently used to describe different classes of interactions depending upon the study. Frequently, studies do not differentiate between *antagonistic* and *reversal* interactions simply classifying both as being *antagonistic*.

**Supplementary Material 5** – *Weighted Multi-Variate / Multi-Level Meta-Analyses*

Meta-analyses were conducted using the *metafor* package, (Viechtbauer, 2010), in R, using the function *rma.mv()*. The meta-analyses conducted within our research were weighted. For every meta-analyses, each interaction was weighted, with the weighting given to each interaction by determined by the *rma.mv()* function.

Many of the included empirical studies conducted experiments where multiple treatments of the same stressor were investigated, or the same control was used for multiple factorial experiments. In order to account for covariance between these interactions, covariance-variance matrices were used within the empirical meta-analyses. Covariance-variance matrices take the form of the below example, (Equation S5.1). In this example the variance of the four interactions are along the major diagonal, ($v_i$), whilst interactions 1 and 2 share a control, hence the covariance of these two interactions, ($c_{1,2}$), is incorporated in the corresponding off-diagonal positions. Where two interactions do not share a control, (there is no covariance), the corresponding off-diagonal position is filled with a 0.

$$\begin{pmatrix} v_1 & c_{1,2} & 0 & 0 \\ c_{1,2} & v_2 & 0 & 0 \\ 0 & 0 & v_3 & 0 \\ 0 & 0 & 0 & v_4 \end{pmatrix}$$

Equation S5.1, an example of a covariance-variance matrix.

For the empirical meta-analyses, random effects were specified as being:

i) The study the interaction was published in,

ii) The study group of organisms.

Within the meta-analytical model, ii) was nested within i). Accordingly, i) allows us to account for between study variance, whilst ii) allows us to account for within study variance.

For the meta-analytical results within the main text of the paper, there were no fixed effects specified. However, two additional meta-analytical models were conducted, with the categorical moderators of *stressor pair* and *organism group*, to test whether our results were robust across different subsets of the empirical data, (Figures S5.1 and S5.2). The results from these meta-analytical models are in line with the main findings of this study. All sub-groups reported summary effect sizes that were negative, with confidence that did not overlap zero, (i.e. antagonistic/reversal interactions), or had confidence intervals that overlapped zero, (additive interactions). However, it should be noted that many of these sub-groups had small numbers of interactions, and as such care should be taken when inferring conclusions from these results.

For the theoretical meta-analytical models, a different approach was used. This was due to computational constraints. Instead, weighted fixed effect meta-analytical models were conducted using the *lm* function in R. For models using the theoretical data, the weighting given to each interaction was specified as being the reciprocal of the interaction variance. Discrepancies between the use of *metafor*, *lm*, *nlme*, and *lme4* have been previously documented, (see <http://www.metafor-project.org/doku.php/tips:i2_multilevel_multivariate>). Ultimately, it has been shown that the summary effect size between the two methodologies remains the same, but that the standard errors differ. While we acknowledge the differences between the two methodologies, due to the computational constraints it was unavoidable.

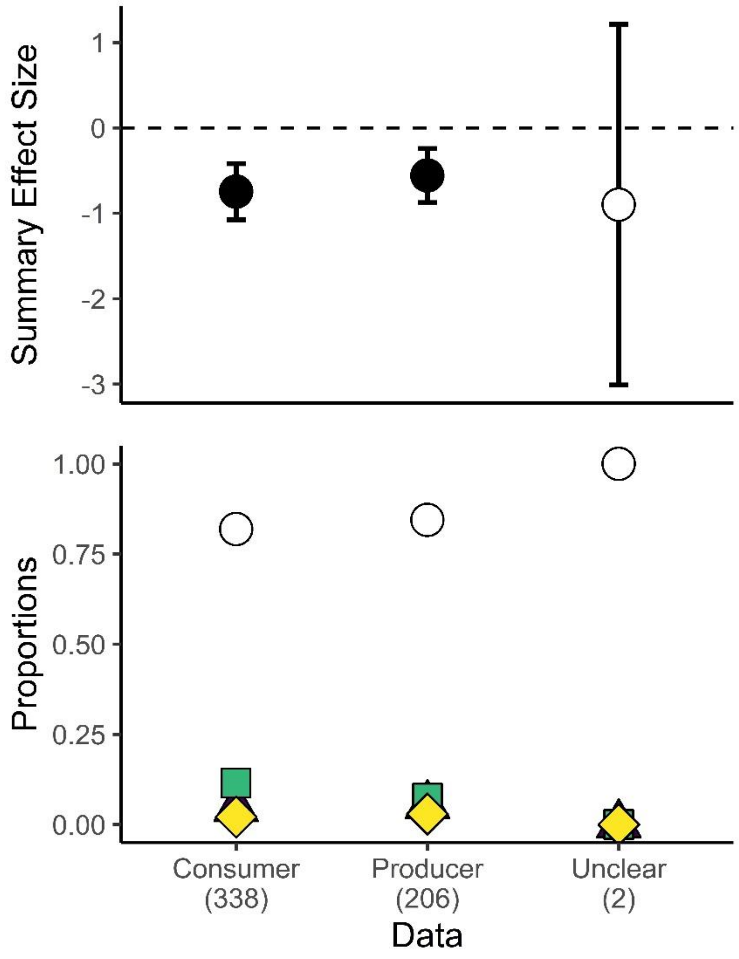


Figure S5.1 Top Panel: Results of meta-analytical model with the inclusion of organism group, (i.e. producer, consumer, or unclear), as a fixed effect. For each organism group, the summary effect size, with accompanying 95% confidence intervals are shown. Filled circles indicate that the 95% confidence intervals do not overlap zero, while filled circles indicate that the 95% confidence intervals do overlap zero. Bottom Panel: the proportions of the different interaction classes, for each organism group, legend as for Figure 3. For x-axis labels, numbers in brackets indicate the number of interactions for organism group.

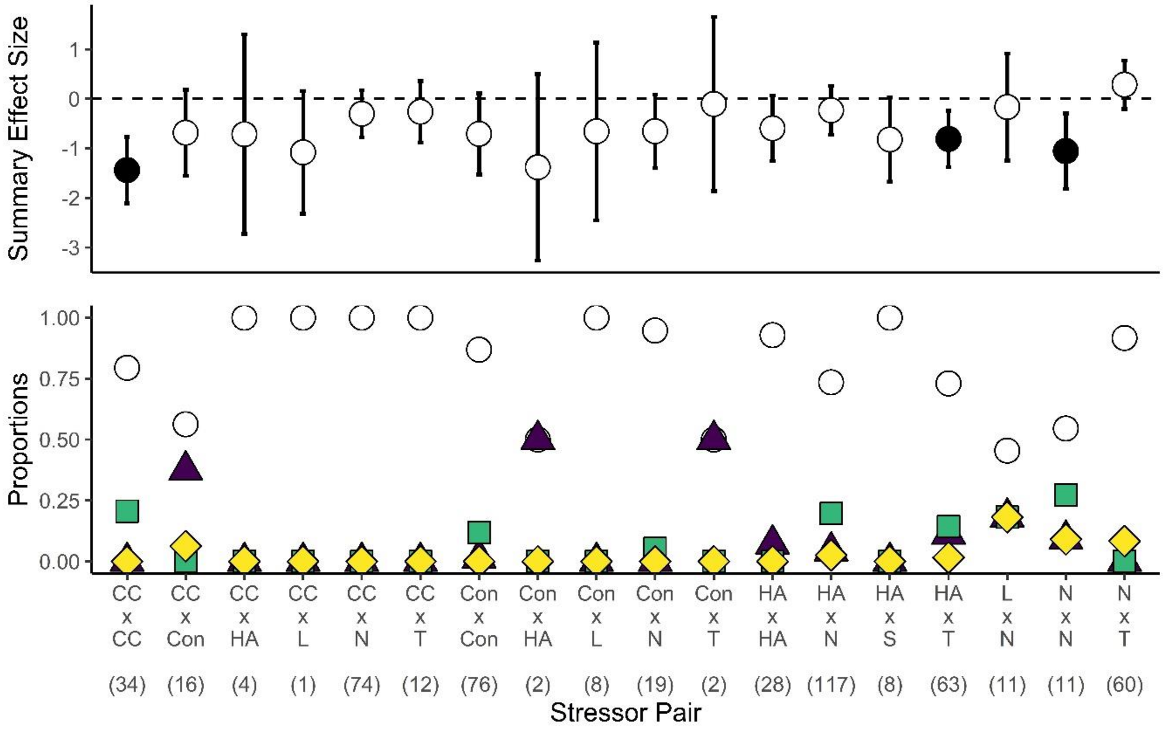


Figure S5.2 Top Panel: Results of meta-analytical model with the inclusion of stressor pair as a fixed effect. For each combination of stressors, the summary effect size, with accompanying 95% confidence intervals are shown. Filled circles indicate that the 95% confidence intervals do not overlap zero, while filled circles indicate that the 95% confidence intervals do overlap zero. Bottom Panel: the proportions of the different interaction classes, for each pair of stressors, legend as for Figure 3. For x-axis labels, numbers in brackets indicate the number of interactions for each pair of stressors. The various acronyms are defined as follows: *CC – Community Composition, Con – Contamination, HA – Habitat Alteration, L – Light, N – Nutrients, T – Temperature.*

Within the main text of the paper, when comparing between empirical and theoretical interactions, (Figure 3), only theoretical interactions with an observation error of $\geq 1x10^{-2}$ were used. This level of observation error was selected as it is likely to represent ecologically relevant levels of observation error. Additionally, in the below figures, (Figures S5.3-S5.5), we show how the proportions of interactions, and summary effect sizes, vary when different ranges of observation error are used. As expected, by expanding the range of observation errors considered, the predictions deviate from those seen in the empirical data.

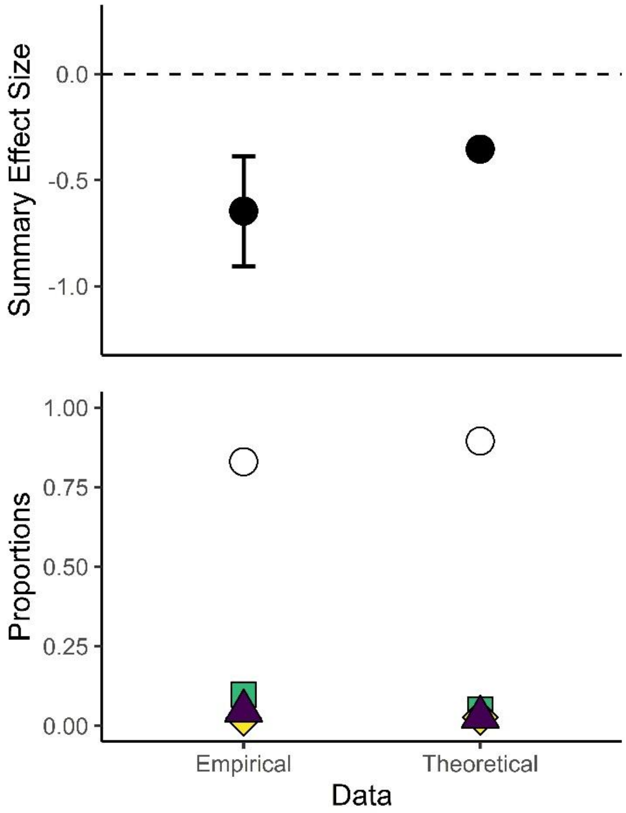


*Figure S5.3: Summary effect sizes, and proportions of the different interaction classes, for the empirical, and theoretical dataset. The empirical dataset comprised 546 interactions, while the theoretical dataset comprised all interactions using observation errors between $1x10^{-1}$ and 0.5. White circles denote additive interactions. Green squares denote antagonistic interactions. Yellow diamonds denote synergistic interactions. Purple triangles denote reversal interactions.*

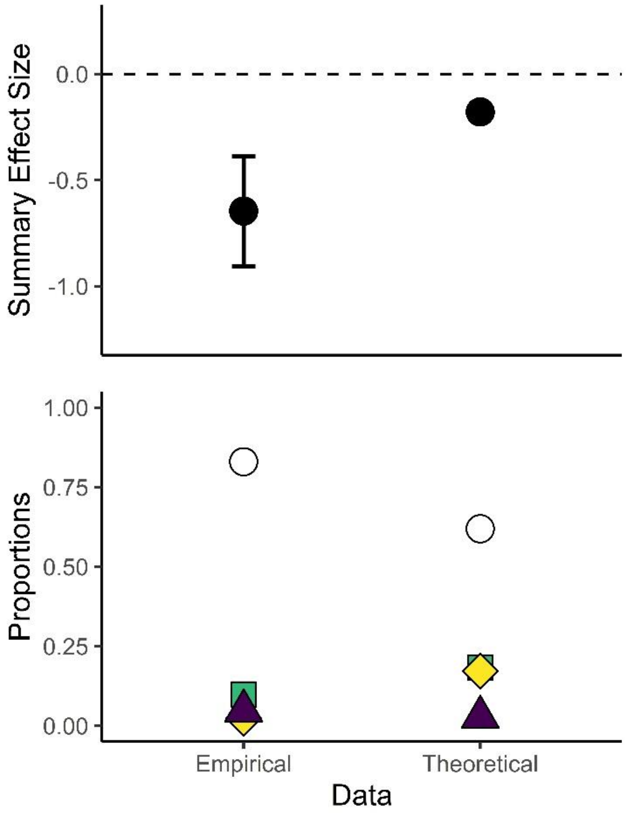


*Figure S5.4: Summary effect sizes, and proportions of the different interaction classes, for the empirical, and theoretical dataset. The empirical dataset comprised 546 interactions, while the theoretical dataset comprised all interactions using observation errors between 1x10$^{-3}$ and 0.5. White circles denote additive interactions. Green squares denote antagonistic interactions. Yellow diamonds denote synergistic interactions. Purple triangles denote reversal interactions.*

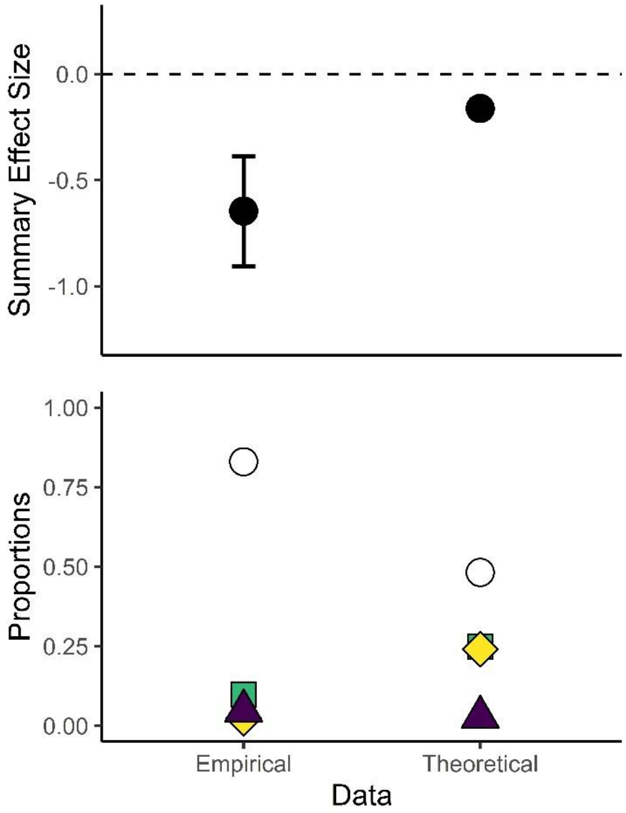


*Figure S5.5: Summary effect sizes, and proportions of the different interaction classes, for the empirical, and theoretical dataset. The empirical dataset comprised 546 interactions, while the theoretical dataset comprised all interactions using observation errors between $1x10^{-4}$ and 0.5. White circles denote additive interactions. Green squares denote antagonistic interactions. Yellow diamonds denote synergistic interactions. Purple triangles denote reversal interactions.*

**Supplementary Material 6** – *Meta-Analysis Heterogeneity and Asymmetry*

*Heterogeneity*

In order to quantify the effects of heterogeneity within the meta-analytical models, $I^2$ statistics were calculated. For the overall meta-analytical model an $I^2$ statistic of 47.0%, just below the benchmark of high heterogeneity from Higgins et al. (2003); though the usefulness of these benchmarks in ecological meta-analyses is questioned, (Nakagawa et al., 2017). Overall, this suggests that heterogeneity within our meta-analysis was lower than either the mean, (91.69%), or median, (84.67%), $I^2$ found in an analysis of previous ecological meta-analyses, (Senior et al., 2016).

Subsequently, in order to explore the some of the sources of heterogeneity we conducted two additional meta-analyses on subgroups of the empirical dataset using the ecologically relevant categorical moderator of organism group, (namely Producer or Consumer), to subset the data. Overall there were 206 interactions for producers, and 338 interactions for consumers, (while a further two interactions were classed as *unclear* – namely where the response of dinoflagellates to interacting stressors was measured – and excluded from further analysis). Overall, running meta-analytical models following the same methodology as outlined in the methods, the meta-analytical model for producers returned an effect size of -0.573 ±0.384, corresponding to an additive summary interaction class; while the meta-analytical model for consumers returned an effect size of -0.629 ±0.331, corresponding to a summary interaction class of antagonistic/reversal. For both of these sub-group meta-analyses $I^2$ statistics were calculated. For the producer meta-analytical model, $I^2$ was equal to 67.8%, while for the consumer meta-analytical model, $I^2$ was equal to 40.5%. Overall, the sub-group analysis revealed that even within these sub-groups, heterogeneity was either moderate or high.

*Publication Bias*

Within the overall meta-analysis, moderate-high levels of heterogeneity were observed. However, this can have ramifications for the traditional methods of assessing publication bias within meta-analyses,

(Ioannidis & Trikalinos, 2007; Nakagawa et al., 2017). Publication bias can manifest through a lack of studies where there are small, or insignificant, effects being published, and instead studies reporting large, or significant effects, are predominately published. As has been previously noted, ecological meta-analyses frequently report high levels of heterogeneity (Senior et al. 2016), as such assessing publication bias can be challenging, as indications of publication bias may instead reflect high levels of heterogeneity.

For our analysis we report the Fail-Safe Number, (using the Rosenberg approach), which indicates the number of studies averaging null results that would need to be added to the analysis to reduce the significance level to a target alpha level, (0.05). For our analysis the reported Fail-Safe Number is 33674.

Additionally, we include two funnel plots illustrating the relationship between effect size and sample size, (Figure S6.1a) and effect size and the inverse of the interaction variance, (S6.1b). As shown by Figure S6.1a, there are consistent bands of sample sizes which correspond to sample sizes exactly devisable by four, (i.e. 8, 12, 16, 20, 32), which correspond to equal numbers of replicates for each of the four treatments within each interaction. Similarly, these bands are also visible within Figure S6.1b. Interpretation of these plots is difficult for several reasons. Firstly, there is correlation between both sample sizes and effect sizes, and the variance and the effect sizes. This is because the sample size is used in the calculation of the effect size and the effect size is used in the calculation of the variance (See Supplementary Material 3). Further complicating this is the complex structure of our data, and the inability of the plots to accurately reflect this. While the meta-analytical models account for both covariance between interactions where interactions share a control or control and treatment (See Supplementary Material 5), or the specified random effects (See Supplementary Material 5), this is not able to be accounted for within these plots. Similarly, we implement Egger's test, (modified to account for multivariate/multi-level meta-analyses), which is a measure of funnel plot symmetry, using the inverse of interaction variance as the specified moderator, (p-value < 0.001) which indicated significant asymmetry within the funnel plot. However, again we note both the potential difficulties in interpreting funnel plots, and the difficulties which heterogeneity poses to identifying publication bias.

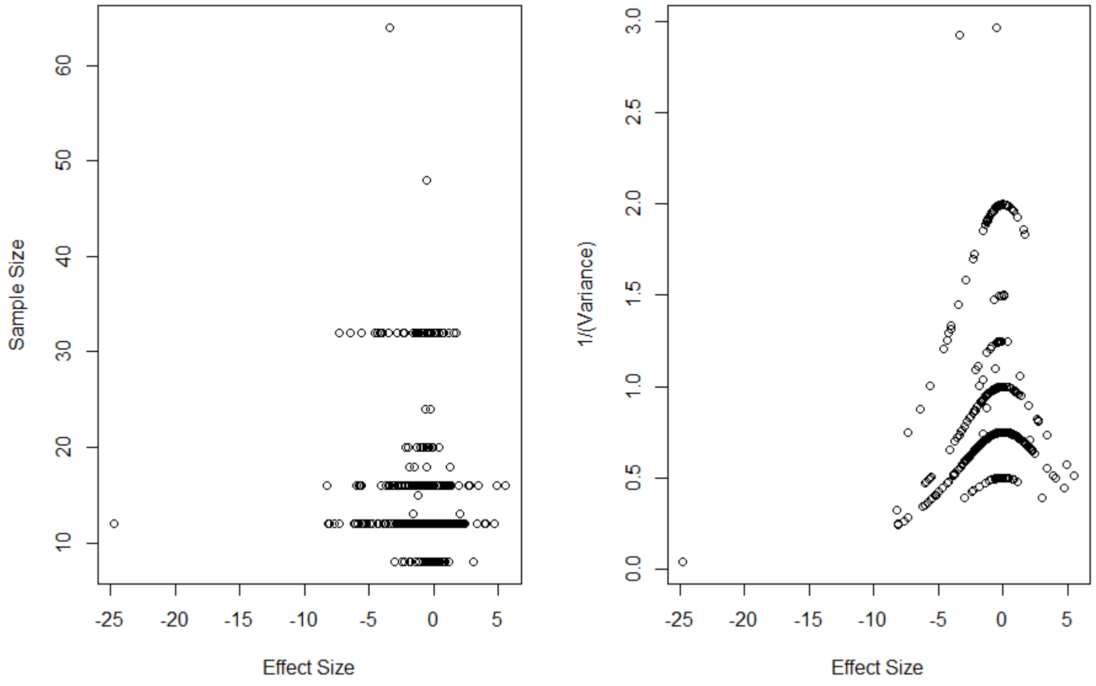


Figure S6.1: a – Sample size against interaction effect sizes; b – inverse of the interaction variance against interaction effect size.

**Supplementary Material 7** – *Empirical Data Reference List*

Below is the reference list for studies from which empirical data was extracted, (listed in alphabetical order).

9. Cottingham, K.L., Glaholt, S. and Brown, A.C., 2004. Zooplankton community structure affects how phytoplankton respond to nutrient pulses. *Ecology*, 85(1), pp.158-171.
10. Dahl, J., 1998. The impact of vertebrate and invertebrate predators on a stream benthic community. *Oecologia*, 117(1-2), pp.217-226.
11. Dalton, R.L., Boutin, C. and Pick, F.R., 2015. Determining in situ periphyton community responses to nutrient and atrazine gradients via pigment analysis. *Science of the Total Environment*, 515, pp.70-82.
12. Davis, S.J., Mellander, P.E., Kelly, A.M., Matthaei, C.D., Piggott, J.J. and Kelly-Quinn, M., 2018. Multiple-stressor effects of sediment, phosphorus and nitrogen on stream macroinvertebrate communities. *Science of the Total Environment*, 637, pp.577-587.
13. De Stefano, L.G., Gattás, F., Vinocur, A., Cristos, D., Rojas, D., Cataldo, D. and Pizarro, H., 2018. Comparative impact of two glyphosate-based formulations in interaction with *Limnoperna fortunei* on freshwater phytoplankton. *Ecological Indicators*, *85*, pp.575-584.
14. Duarte, S., Pascoal, C., Alves, A., Correia, A. and Cassio, F., 2008. Copper and zinc mixtures induce shifts in microbial communities and reduce leaf litter decomposition in streams. *Freshwater Biology*, 53(1), pp.91-101.
15. Fernández-Aláez, M., Fernández-Aláez, C., Bécares, E., Valentín, M., Goma, J. and Castrillo, P., 2004. A 2-year experimental study on nutrient and predator influences on food web constituents in a shallow lake of north-west Spain. *Freshwater Biology*, 49(12), pp.1574-1592.
16. Ferragut, C. and de Campos Bicudo, D., 2012. Effect of N and P enrichment on periphytic algal community succession in a tropical oligotrophic reservoir. *Limnology*, *13*(1), pp.131-141.
17. Gardeström, J., Ermold, M., Goedkoop, W. and McKie, B.G., 2016. Disturbance history influences stressor impacts: effects of a fungicide and nutrients on microbial diversity and litter decomposition. *Freshwater Biology*, *61*(12), pp.2171-2184.
18. Gattás, F., De Stefano, L.G., Vinocur, A., Bordet, F., Espinosa, M.S., Pizarro, H. and Cataldo, D., 2018. Impact of interaction between Limnoperna fortunei and Roundup Max® on freshwater

phytoplankton: An in situ approach in Salto Grande reservoir (Argentina). *Chemosphere*, 209, pp.748-757.

**Supplementary Material 8** – *Summary Figures Empirical Data*

Additional figures detailing the geographic distribution of papers from which empirical data was extracted, (Figure S8.1), and the number of papers per year that were included in this analysis, and the number of papers per year that were revealed in the systemic literature search, (Figure S8.2).

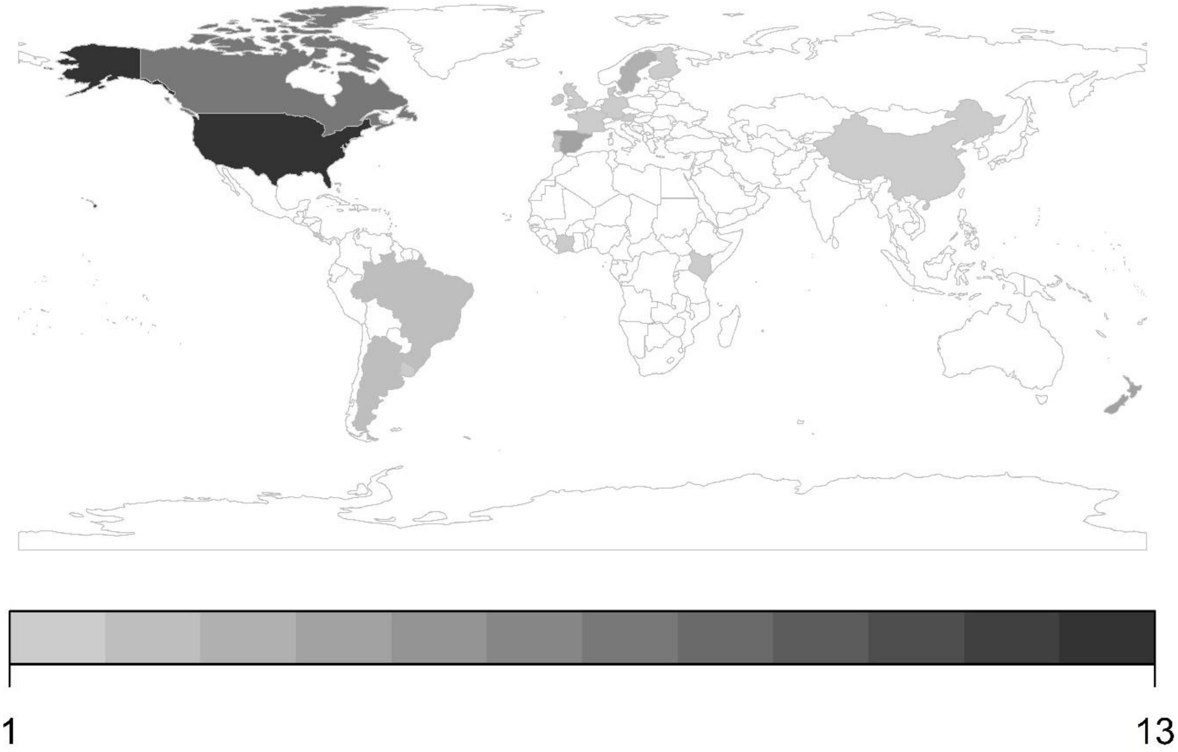


Figure S8.1: Number of papers, from which data was extracted, that were included in our analysis shown by country of origin. Darker colours indicate more papers originating from the country in question. Countries coloured white, indicate no papers originating from the country in question.

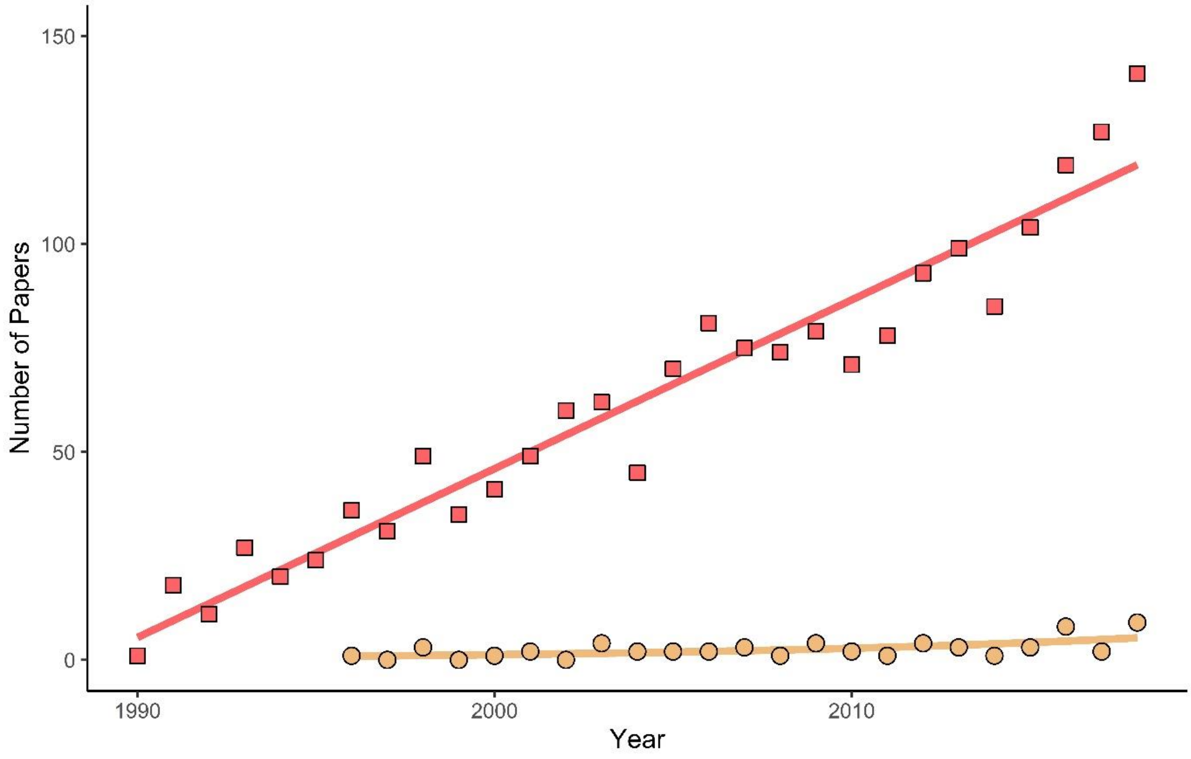


Figure S8.2: Number of papers by year that were revealed by the systematic literature review, (red squares), and those that were included in our study, (orange circles). Lines indicate generalised liner models fit to the data, with Year as the explanatory variable and Number of Papers as the response variable, both Gaussian and Poisson distributions were used to fit the models, with AIC values used to select the best fitting model. For the systematic literature review, a Gaussian distribution with intercept = -8074.1961, p<0.001; and coefficient =4.0601, p<0.001 was the best fitting model. For the papers included within our analysis, a Poisson distribution with intercept = -160.25750, p<0.001; and coefficient = 0.08024, p<0.001 was the best fitting model.
